# scLEMBAS: Context-Aware Modeling of Signaling Pathway Activity at Single-Cell Resolution

**DOI:** 10.64898/2026.07.20.739670

**Authors:** Hratch M. Baghdassarian, Nikolaos Meimetis, Olof Nordenstorm, Brian A. Joughin, Avlant Nilsson, Douglas A. Lauffenburger

## Abstract

Cells sense and integrate extracellular cues through intracellular signaling networks that reshape transcription factor activity to dictate cellular responses. Signaling activity is difficult to decipher: it is non-linear, and it contains extensive feedback and crosstalk. Furthermore, the same perturbation can elicit markedly different responses depending on context (e.g., cell type, disease state, and tissue microenvironment) such that identical stimuli produce diverse responses in multicellular populations. Consequently, there is a vast combinatorial space of complex interactions and context-dependent responses that necessitate computational models. Computational models of single-cell perturbation responses are demonstrated to predict cellular responses, but are often limited in mechanistic insight. Prior knowledge networks offer a route to bridge predictive capability and interpretability. Here we present scLEMBAS, a context-aware, gray-box neural network that models signaling pathway activity at single-cell resolution while preserving mechanistic grounding. scLEMBAS encodes a prior-knowledge network of protein–protein interactions as a recurrent neural network whose learnable edge weights correspond to signaling interaction strengths. It also captures context and individual cell variance through compositional bias terms. An adversarial approach allows the model to answer a single-cell counterfactual – what a given cell’s TF activity would be under a different perturbation or context – while involving mechanistic rather than simply relational information. Across two scRNA-seq datasets spanning single- and multi-perturbation settings, scLEMBAS accurately predicts out-of-distribution combinations of perturbation and context. Capturing population variance across individual cells enables the model to predict cell subtype specific perturbation responses, despite being agnostic to such labels. Beyond prediction, scLEMBAS’ learned parameters are biologically interpretable: learned edge weights carry information beyond network topology and "self-prune" spurious interactions, while the categorical bias nominates proteins associated with cell-type-specific perturbation states. Overall, scLEMBAS enables quantitative dissection of how signaling pathway activity is reshaped by perturbation within specific cellular contexts.

## Introduction

Cells sense and integrate extracellular cues through intracellular signaling networks that alter transcription factor (TF) activity, ultimately affecting cellular phenotype. Rather than proceeding along isolated, linear paths from receptor to TF, the signal is propagated through a densely interconnected network characterized by non-linearity^1^, feedback, and pathway crosstalk^2^. Such properties enable the cell to compute complex responses to multiple stimuli^3^. Critically, the same stimulus can elicit markedly different responses depending on context—the cell type, its state, and its local microenvironment—such that identical perturbations produce heterogeneous outcomes across a population^4^. Single-cell RNA sequencing (scRNA-seq) has rendered this heterogeneity directly observable, and, combined with multiplexed perturbation screens, now permits systematic interrogation of how distinct cell populations respond to defined stimuli^5^.

The aforementioned signaling network properties (e.g., non-linearity, crosstalk) make pathway activity difficult to decipher. Furthermore, the vast combinatorial space of possible interactions^3^, perturbations, and contexts cannot be exhaustively surveyed experimentally. Consequently, computational models are necessary to predict and interpret signaling responses across this space. A growing family of machine-learning methods addresses this need by learning to predict single-cell perturbation responses; one common such approach is the decomposition of measured variation into additive contributions from perturbation, cell type, and other covariates in order to extrapolate to unseen combinations (e.g., compositional perturbation autoencoder^6^). Such models can predict responses to held-out dosages, cell types, and drug combinations, and have proven valuable for *in silico* hypothesis generation^7^. However, they are often statistical in nature: they capture co-expression programs correlated with a perturbation and, due to the “black-box” nature of neural networks, their learned representations bear no explicit correspondence to the molecular circuitry of the cell. Consequently, predictions are difficult to attribute to specific mechanisms, and the models offer limited insight into which signaling components might be tuned to redirect a response.

Prior knowledge offers a route to bridge the gap between the predictive capability of complex responses and mechanistic insight^7,8^. The topology of the signaling network has been extensively catalogued in curated resources of protein–protein interactions (PPIs), such as OmniPath^9^, and embedding this topology directly into a model constrains its parameters to biologically meaningful interactions. We previously developed such an approach, LEMBAS^10^, for modeling bulk signaling pathway activity that exemplifies this strategy: it encodes a prior knowledge network (PKN) as a recurrent neural network (RNN) whose learnable weights correspond to the strengths of individual edges, taking ligand concentration (or intracellular signaling pathway perturbations) as input and TF activity as output. Because the architecture is derived from the signaling topology, LEMBAS provides a means for genome-scale signaling pathway modeling that can capture feedback (specifically captured by the RNN) and non-linearity while preserving mechanistic interpretability: its parameters can be read as quantitative signaling interactions^11^, and it supports functionalities such as *in silico* gene knockouts and identification of off-target drug effects^12^. The PKN additionally acts as a strong structural prior—the sparsity it enforces reduces the effective parameter space. These properties make PKN-constrained models particularly well suited to extracting mechanism from noisy, high-dimensional data^7^. However, because LEMBAS was developed for bulk measurements, it cannot discriminate between cell types without additional parameterization from gene expression, generalize across the multiple contexts that shape a response, or capture the cell-to-cell dispersion intrinsic to single-cell data.

Here we present scLEMBAS, a context-aware extension of LEMBAS that models signaling pathway activity at single-cell resolution while preserving its mechanistic grounding. scLEMBAS retains the PKN-constrained RNN but decomposes the LEMBAS’ bias term, which represents protein-specific activity, into two additive, conceptually distinct components: a categorical bias that captures context-specific activity such as cell type and *a global bias* that captures the residual heterogeneity beyond context and perturbation (Fig. 1A). This factorization allows scLEMBAS to answer a counterfactual—what a given cell’s TF activity would be under a different perturbation or context. In doing so, scLEMBAS unites the interpretability of mechanistic, prior-knowledge-based models with the flexibility of statistical single-cell perturbation predictors.

**Fig. 1:**
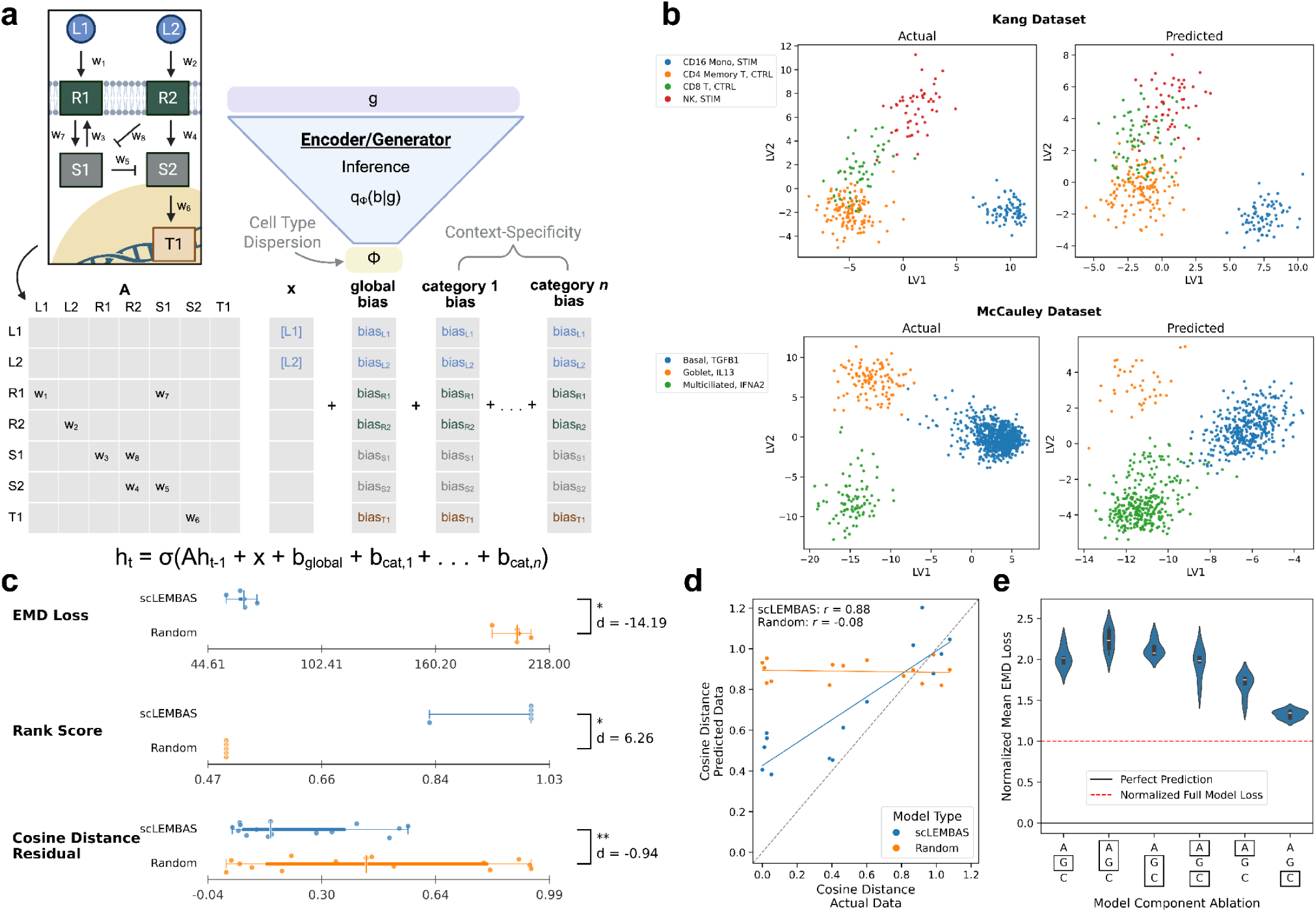
scLEMBAS Architecture Accurately Captures OOD TF Activity Across Multiple Metrics. **(a)** An adjacency matrix “**A**” representing the signaling network interactions is combined with the perturbation inputs “**x**” and bias term. All parameters are input to a Michaelis-Menten activation function and adjusted by a recurrent neural network (RNN). To capture single-cell heterogeneity, a stochastic encoder can output the global bias, which is devoid of perturbation- and context-information. This is generated from the gene expression “**g**”. For context-specificity, the bias term is decomposed into the global bias (baseline activity) and categorical terms (context-specific activity). **(b)** Scatter plots of TF activity of OOD representative test-split OOD conditions in latent space. Latent embeddings were fitted using the observed data (left panels), after which predictions were projected into the learned space (right panels). The latent space was defined by a PLS-reduced space using the test condition as the response variable. **(c)** scLEMBAS’ predictive performance assessed by three different metrics. Individual test conditions (cosine residual metric) or condition-wise means within folds (EMD loss and rank score metrics) are shown as points (with vertical jitter for visual clarity), with the median indicated by a vertical line, the interquartile range (25th–75th percentile) by the thick bar, and the full range (minimum to maximum) by the thin whisker. Models are compared against a random baseline trained on permuted features using a paired Wilcoxon signed-rank test (*** p ≤ 0.001, ** p ≤ 0.01, * p ≤ 0.05) and are annotated by a paired Cohen’s d effect size. Each metric is plotted on its own native scale. See Methods for details on metric computations. The cosine distance residual is defined as the absolute distance of each scatter point (test condition) in panel (c) to the identity line, representing deviation from perfect prediction. **(d)** Scatter plots and regression lines compare cosine distances derived from predicted and actual data, with Pearson correlations annotated. Each point corresponds to a unique perturbation in the test split aggregated across the five cross-validation folds. The identity line (gray dashed line) indicates perfect agreement between predicted and observed perturbation geometry. Cosine distances were computed following Viñas Torné et al.^18^ (see Methods for details) and quantify perturbation-specific directional variation relative to shared control effects. **(e)** Violin plots showing the across-fold distribution of normalized EMD loss for models with different component ablations. Within each fold, EMD loss is computed for each test condition and averaged across conditions. For each ablated model forward pass, losses are normalized to the corresponding full scLEMBAS model (red dashed line), such that values greater than one indicate worsened predictive performance (i.e., a positive contribution of the omitted component to model accuracy). X-axis labels denote the ablated component with a black box, with unboxed components retained in the forward pass. “A” represents the adjacency matrix, “G” the global bias, and “C” the categorical bias. Panels (c-e) display results specific to the McCauley dataset, with analogous Kang dataset results visualized in the Supplementary Materials.

We show that scLEMBAS accurately predicts out-of-distribution (OOD; unseen combinations of perturbation and context) TF activity on two scRNA-seq datasets spanning binary and multi-perturbation settings across complementary distributional, rank-based, and geometric metrics. Through targeted ablations, we confirm that each architectural component contributes to predictive performance and behaves in its intended role. The global bias enables counterfactual perturbation prediction while retaining cell subtype-specific variation. The learned PKN edge weight magnitudes carry biological information beyond simple binary network topology (i.e., presence or absence), and that the model “self-prunes” spurious interactions irrelevant to the biological context of interest. Finally, we demonstrate that the categorical bias can be used to prioritize the signaling proteins defining cell-type-specific perturbation responses—nominating candidates whose *in silico* alteration shifts one cell type’s predicted state toward another’s. Together, these results establish scLEMBAS as a mechanistic, interpretable framework for dissecting context-dependent signaling from single-cell data.

## Results

### A gray box neural network approach to model signaling pathway activity

LEMBAS^10^ encodes a PKN as a RNN, taking signaling perturbations as input and predicting TF activity, inferred from gene expression^13^, as output. We modified and extended the LEMBAS architecture to (1) generalize across multiple contexts (e.g., cell type or disease state) and (2) capture the heterogeneity inherent to single-cell measurements (i.e., variance of individual cells) (Fig. 1A). Each architectural component in scLEMBAS has a specific conceptual grounding in mechanism. The adjacency matrix is constructed on the topology of the PKN of protein-protein interactions (here, Omnipath^9^), enabling the model to globally learn previously incompletely annotated quantitative values for edge weights in response to perturbation. The bias term represents protein-specific activity. Thus, the adjacency matrix represents context-independent interaction strengths between proteins, whereas the bias terms represent protein-specific activity thresholds that modulate signaling in a context-dependent manner. Alongside the perturbation input, these components are combined and non-linearly transformed using a Michaelis-Menten-like activation function within a RNN, enabling the model to capture feedback loops.

To address the goals of context generalization and capturing single-cell heterogeneity, we decomposed the bias term into two additive components: a categorical bias term and a global bias term. To do so, we draw on compositional and adversarial approaches from single-cell perturbation modeling^6,7^. The categorical bias, encoded by learned embeddings, captures context-specific activity (in this case cell type). The global bias, generated by a stochastic encoder from an input gene expression vector, captures single-cell heterogeneity. Although gene expression contains the cell-state information needed to capture this heterogeneity, providing it directly would allow the model to bypass learning the effects of perturbation and categorical context, thereby limiting OOD generalization. We therefore implement adversarial removal to retain residual cell-state variation while removing perturbation- and context-associated information.

The predictive task of scLEMBAS is to predict OOD conditions (Fig. S1): unseen combinations of cell type and perturbation, but where each individual cell type and perturbation is observed during training. Through the adversarial removal approach, scLEMBAS enables counterfactual prediction of the TF activity of a given cell under a different condition.

### scLEMBAS Accurately Predicts Out-of-Distribution Transcription Factor Activity

We assessed scLEMBAS on two single-cell RNA-sequencing datasets: the “Kang” dataset representing peripheral blood mononuclear cells (PBMCs) in a binary perturbation setting (unstimulated or stimulated with a fixed concentration of 100 U/mL IFNβ)^14^ and the “McCauley” dataset representing the airway epithelium under multiple perturbations representing distinct downstream signaling activation^15^ (Fig. S2; Table 1). For each dataset, sets of individual combinations of cell type and perturbations, (e.g., stimulated CD16 monocytes in Kang) were removed from the data during model training across a 5-fold cross-validation splitting procedure, and their TF activity levels predicted from the remaining data (see Methods for details). Given that the field has not converged on a single definitive metric for evaluating single-cell perturbation predictions (see Supplementary Information and Discussion for details)^16,17^, we first conduct a visual inspection of predictions in latent space (Fig. 1b), enabling qualitative assessment of model behavior, and subsequently employ a diverse range of quantitative metrics (Fig. 1c). Specifically, we quantified performance using three complementary metrics capturing global, relative, and geometrical aspects of prediction: (i) the mean per-condition Earth Mover’s Distance (Wasserstein-1), which measures distributional similarity between predicted and observed single-cell responses, (ii) a rank-based metric introduced by Wu et al.^17^, which evaluates the relative position of each perturbation prediction relative to all other predicted perturbations, and (iii) a cosine distance based metric derived from Viñas Torné et al.^18^, which captures the relative geometry of specific perturbations relative to the control and average perturbation effect.

Visually, we observe that across folds, scLEMBAS’ predictions largely preserve the relative geometry of held-out test conditions, as exemplified by representative test condition latent space visualizations (Fig. 1b). This qualitative observation is supported by quantitative comparisons to a random baseline model fit on permuted features. Specifically, scLEMBAS exhibits substantially higher agreement between predicted and observed cosine distances relative to random predictions (Fig. 1c-d). The cosine distance quantifies the specificity of a given perturbation relative to all other perturbations^18^. As such, the high Pearson correlation (*r* = 0.88), together with the low distance to the identity line, indicates that scLEMBAS captures both systemic perturbation effects and perturbation-specific directional structure present in the actual data; the relatively small effect size of the residual distance compared to other metrics is likely reflects the random baseline’s lack of correlation with the actual data (Fig. 1d, r = −0.08), which produces a near-horizontal relationship and artificially small distances to the identity line at high values. scLEMBAS predictions also perform significantly better than random as quantified by the Earth Mover’s Distance (EMD) loss and rank-based metric introduced by Wu et al.^17^ (Fig. 1c, Fig. S8a). While EMD represents a commonly used single-cell-resolution predictive performance metric, the rank-based score has been proposed as a complementary measure that emphasizes the relative perturbation specificity of predictions^17^. The random model ranks tend to degenerate to 0.5, the expected score of random predictions.

In addition to comparisons against a random baseline, we also compared scLEMBAS’ predictive performance against pseudo-bulked models (random forest, linear, and training-mean baselines) described previously^19,20^, which are intended to represent deliberately simple baselines of capturing perturbation effects. These comparisons showed that evaluating scLEMBAS with the relevant biological context improved the relative performance: scLEMBAS was competitive with or outperformed baselines when evaluation was conducted in latent spaces reducing the TF features to the primary axes of variation of interest (e.g., perturbation effects), particularly when cell-type context was considered (Fig. S9, S10, S11). For additional details of the pseudobulk baseline comparisons see the Supplementary Materials.

Importantly, scLEMBAS’ utility extends beyond predictive performance alone, as its ability to learn signaling-network topology and capture specific network nodes involved in perturbation responses provides mechanistic insight unavailable to other models; this will be discussed extensively in the subsequent sections. Given this mechanistic nature, we sought to verify that each of the three aforementioned model components (the adjacency matrix, categorical bias, and global bias) contributes meaningfully and in its intended role. To assess whether each component contributes meaningfully, we performed targeted ablations and quantified the resulting relative change in predictive performance. Across all folds, ablation of any component (i.e., exclusion of that component during prediction) led to degraded performance, indicating that each contributes usefully to model prediction (Fig. 1e, Fig. S8b-c). Consistent with this, ablation of pairs of components tended to have a larger performance degradation than performance of a single component. In the proceeding sections, we further probe the conceptual and functional contributions of these individual architectural components.

### Global Bias Enables Counterfactual Perturbation Prediction

By removing cell type and perturbation information from the encoding of gene expression, the global bias enables appropriate counterfactual prediction. We first verified that this information was effectively removed from the global bias. To do so, we applied our standard principal component analysis (PCA) pipeline (see Methods for details) to the generated bias terms in the test splits. We compared results to a baseline model that did not receive adversarial penalization during training, such that it did not have pressure to remove this information. Visual inspection of the PCA space indicated the reduction of perturbation and cell type information in the global bias relative to this baseline, while inclusion of the categorical bias reintroduces cell type structure (Fig. 2a, Fig. S12). To rigorously assess removal of both linear and nonlinear structure, we trained logistic regression and random forest probing classifiers^21^ to predict cell type and perturbation labels from the generated bias. Adversarial training in scLEMBAS led to significantly lower probe accuracy compared to the baseline model without adversarial penalization for both cell type and perturbation across all folds, with the exception of incomplete removal of linear information in the third fold of the McCauley dataset (Fig. 2b, Fig. S12). Although probing classifier accuracy suggests some residual categorical information in the global bias relative to random chance, we next confirm that information removal is sufficient to enable counterfactual prediction.

**Figure 2:**
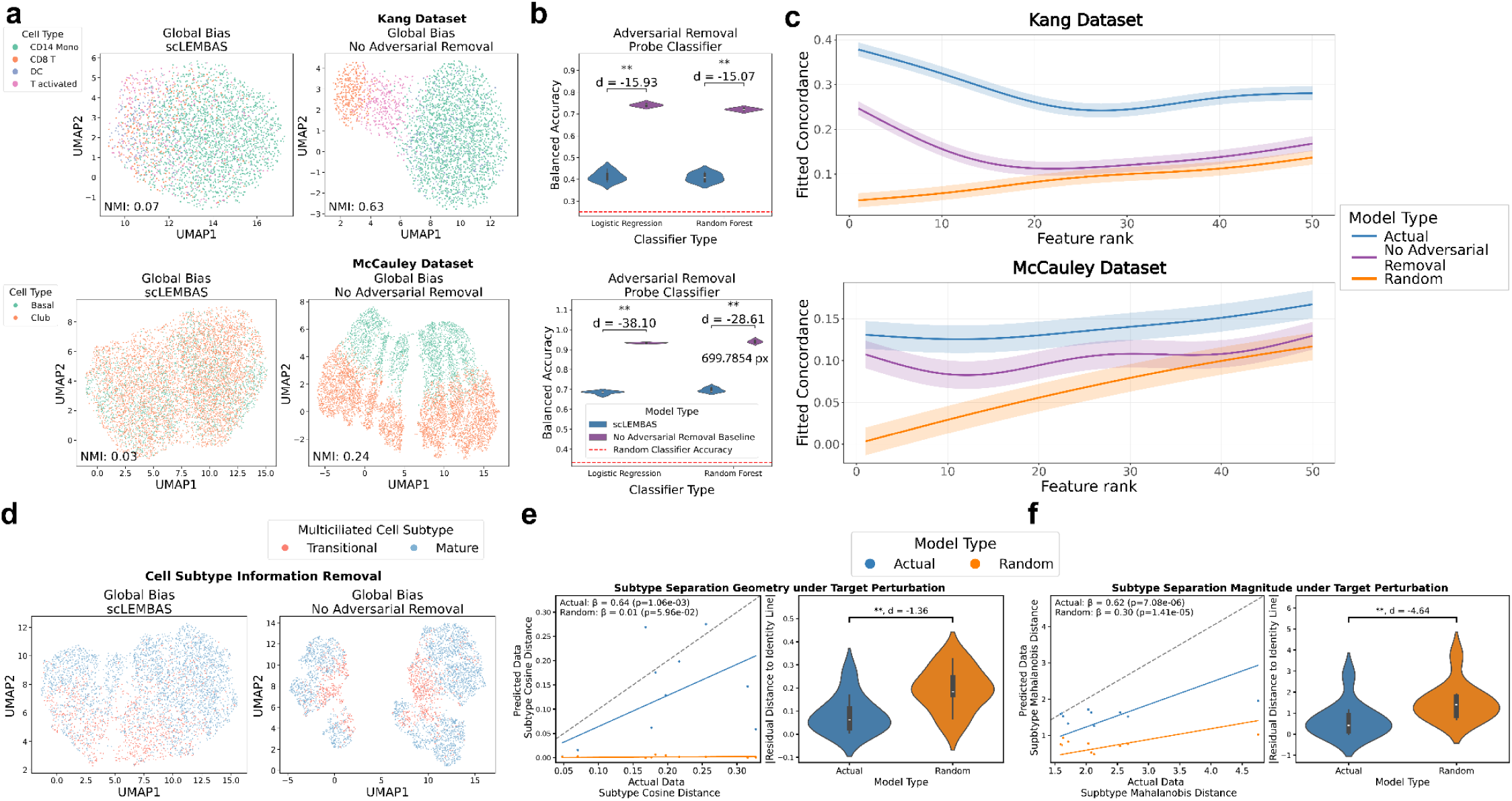
Adversarial information removal enables counterfactual prediction while retaining residual variance in the global bias. **(a)** UMAPs are shown for a representative test-split OOD condition. UMAPS are generated using our standard PCA embedding pipeline on generated bias terms and are colored by cell type. The left panel shows global bias generated from the standard scLEMBAS model in a representative test-split OOD condition. The right panel shows the global bias from the baseline model without adversarial penalization. Panels are annotated with the normalized mutual information between the label of interest and the Leiden cluster. **(b)** Violin plots for the same representative test condition as in panel (a) compare probing classifier 5-fold CV balanced accuracy between the global bias output by scLEMBAS and the baseline model without adversarial removal. Statistical significance is indicated by the Benjamini–Hochberg FDR-corrected one-sided Mann–Whitney U test (** 0.005 < q ≤ 0.01). probing classifiers were trained to predict the target label using features from each bias’s principal component space. The red dashed line indicates the expected balanced accuracy of a random classifier. **(c)** GAMM-estimated partial effects of scLEMBAS and baseline models on differential expression concordance, marginalized over effect-size direction (up- or down-regulated features). Shaded regions represent 95% confidence intervals. Higher concordance indicates better agreement between model predicted and actual data DE features. **(d)** UMAPs are shown for a representative test-split OOD condition and cell type (here, multiciliated cells). UMAPS are generated using our standard PCA embedding pipeline on generated bias terms stratified by a given cell type and are colored by cell subtype. UMAPS are displayed for the global bias generated by scLEMBAS (left panel) and the global bias generated by the baseline model without adversarial penalization (right panel). **(d)** Scatter plots and OLS regression lines compare cosine distances derived from predicted and actual data (left panel). Each point corresponds to a single counterfactual prediction of cells under the target perturbation given the origin perturbation. Cosine distance is computed between two vectors, each originating at the centroid of the higher-order cell type under the origin perturbation and terminating at the centroids of the two subtypes under the target perturbation; the origin is always the actual data, whereas the target is the model predictions or actual data for the y- and x-axis respectively. OLS coefficients are annotated alongside their Wald test p-values. Coefficients were estimated using models without an intercept to enable direct comparison to the identity line (gray dashed line), with coefficients closer to 1 indicating more accurate preservation of subtype-specific perturbation response geometries. Violin plots summarize the absolute distance of each scatter point to the identity line (right panel), representing deviation from perfect prediction. Comparisons between scLEMBAS and the random baseline model are annotated with a Benjamini–Hochberg FDR-corrected two-sided paired Wilcoxon signed-rank test (** 0.005 < q ≤ 0.01) and a paired Cohen’s d effect size. **(f)** Panels visualize a similar analysis as in panel (e), but using Mahalanobis distance instead of cosine distance. Mahalanobis distances were computed between cell subtypes under the target perturbation.

Given successful adversarial removal of covariates, we next evaluated that this consequently yielded predicted perturbations that are consistently separated from their corresponding controls in latent space, distinct from overall test-condition accuracy. We confirmed accurate counterfactual separation of perturbation predictions globally across all features using the Mahanalobis distance (Fig. S13; see Supplementary Materials for details). Briefly, we observed that scLEMBAS significantly outperformed the baseline, indicative of its capability to learn this counterfactual structure.

However, we also observed a systematic under-estimation of predicted perturbation separation from control as measured across all features using the Mahalanobis distance, likely associated with the failure modes discussed in the Supplementary Results (Supplementary Results - Global Assessment of Counterfactual Perturbation Predictions). Thus, we asked whether predicted perturbations separated strongly enough to capture mechanistically relevant information. Specifically, for each test condition, we ran differential feature (DE) analysis between a predicted perturbation and its respective actual data control. Next, we assessed how well predicted differential expression agreed with actual differential expression using a modified concordance at the top analysis^22,23^, calculating a running Jaccard index concordance between predicted and true DE features across DE rank as ordered by the Cliff’s delta effect size (Fig. 2c). To quantify scLEMBAS’ performance relative to baselines, we fit a generalized additive mixed model (GAMM) regressing concordance on the model type for the top 50 DE features (see Methods for details); smooth terms and fixed effect intercepts for model type were highly significant (all F-tests p < 2e-16). For the McCauley dataset, across all ranks, scLEMBAS had an average 0.07 higher concordance than the random baseline, and 0.04 higher concordance than the baseline with no adversarial removal as calculated by the normalized area under the curve (nAUC), which quantifies the cumulative difference in concordance between models by integrating their contrast across rank; it similarly had 0.19 and 0.14 higher average concordance in the Kang dataset for the two baseline models respectively. Given the stringency of a Jaccard index metric for concordance, this is a substantial improvement in performance over baselines. Furthermore, scLEMBAS outperformed baselines across all feature ranks (Fig. S14). At each rank, this difference was statistically significant, as the 95% Wald confidence interval for the model-based contrast remained strictly positive.

### Global Bias Enables Prediction of Subtype-Specific Transcriptional Differences by Retaining Residual Variance

Because scLEMBAS builds on an algorithm originally designed for bulk omics measurements^10^, we asked whether extending it to single-cell resolution enables retention of subtype-specific heterogeneity in perturbation responses. Specifically, when scLEMBAS is trained with categorical bias terms defined at the level of higher-order cell types, agnostic to more granular subtypes, do its predictions still capture differences in perturbation response across subtypes? The global bias adversarially removes the known, higher-order cell type label, but, in principle, should preserve the residual subtype-level variance since it is not explicitly removed.

In the McCauley dataset, multiciliated and goblet cells were further annotated by cell state as either “transitional” or “mature”. To assess whether the global bias retains subtype specificity, we re-ran the probing classifier pipeline on cell subtypes within each fold. In contrast to the successful adversarial removal of cell type and perturbation labels, we observed no significant differences in balanced accuracy between scLEMBAS and the no-adversarial baseline (Fig. S15b). Several caveats are worth noting. First, the Cohen’s *d* effect size between scLEMBAS and the baseline remains large, which is expected given that higher-order cell type labels contain mutual information with subtype labels, leading to some degree of subtype information removal. However, the average subtype effect size (−5.0) is substantially smaller than that observed for higher-order cell types (−19.1), despite containing fewer labels and thus being an “easier” classification problem. Second, some balanced accuracies for club cells approach random performance, likely due to class imbalance, with only 7.2% of cells labeled as transitional across retained perturbations; in contrast, multiciliated cells consisted of 25.0% transitional cells. Despite these limitations, visualization of the global bias in UMAP space (Fig. 2d, Fig. S15a) indicates that subtype structure is partially retained, at levels comparable to the baseline model, regardless of the relative loss of cell type and perturbation structure. Together, these results provide preliminary evidence that the global bias retains cell subtype information. We next assess whether this retained structure supports subtype-specific perturbation responses in model predictions.

Given the nature of the counterfactual, in which input cell gene expression vectors in an initial perturbation state are transformed to a predicted final perturbation state, we ensured good representation across subtype perturbation shift geometries (see Supplementary Materials for details regarding expression shift geometries). Visually, across representative examples of each perturbation type, the relative geometry of subtype-specific responses appears preserved in scLEMBAS predictions (Fig. S16). To quantify this, we assessed how well both the magnitude and geometry of subtype separation under target perturbations recapitulate the actual data (Fig. 2d-e), confirming scLEMBAS significantly outperforms the random baseline.

### Learned Adjacency Matrix PPI Weights are Meaningful Beyond Network Topology

A major advantage of scLEMBAS is its ability to learn protein-protein interaction weights within the PKN. This can enable identification of interactions that are mechanistically relevant to the biological data being modeled. The model is already substantially constrained due to the sparsity enforced by the PKN network topology. Thus, we wanted to more deeply explore the utility of learning edge weights beyond network topology. In other words, we wanted to assess the mechanistic relevance of weight magnitude in conjunction with other properties of the PKN, as measured in proxy by scLEMBAS’ predictive performance. While we recently demonstrated this behavior in the bulk version of LEMBAS^11^, here we asked whether scLEMBAS retains this capability despite the increased technical noise of scRNA-seq^24^ and the additional architectural complexity of the model.

We first compared learned model predictions to those of a baseline with binarized edge weights representing only the presence or absence of an interaction (see Methods for details). In this setting, the model with learned weights showed higher test prediction performance than the binarized baseline (Fig. 3a), indicating that the learned weights are relevant beyond simple topology. Yet, qualitative comparison of the change in model performance between scLEMBAS and the binarized baseline and that of the random baseline (Fig. 1c, EMD loss) shows that, relatively, the binarized baseline performs substantially better than random. This possibly indicates that PKN-constrained topology alone provides a useful structural prior for perturbation models. We note a caveat here is that, in the binarized baseline, only the adjacency matrix is modified, such that the contribution to predictive capacity may be derived from the bias terms rather than the adjacency matrix topology.

**Figure 3:**
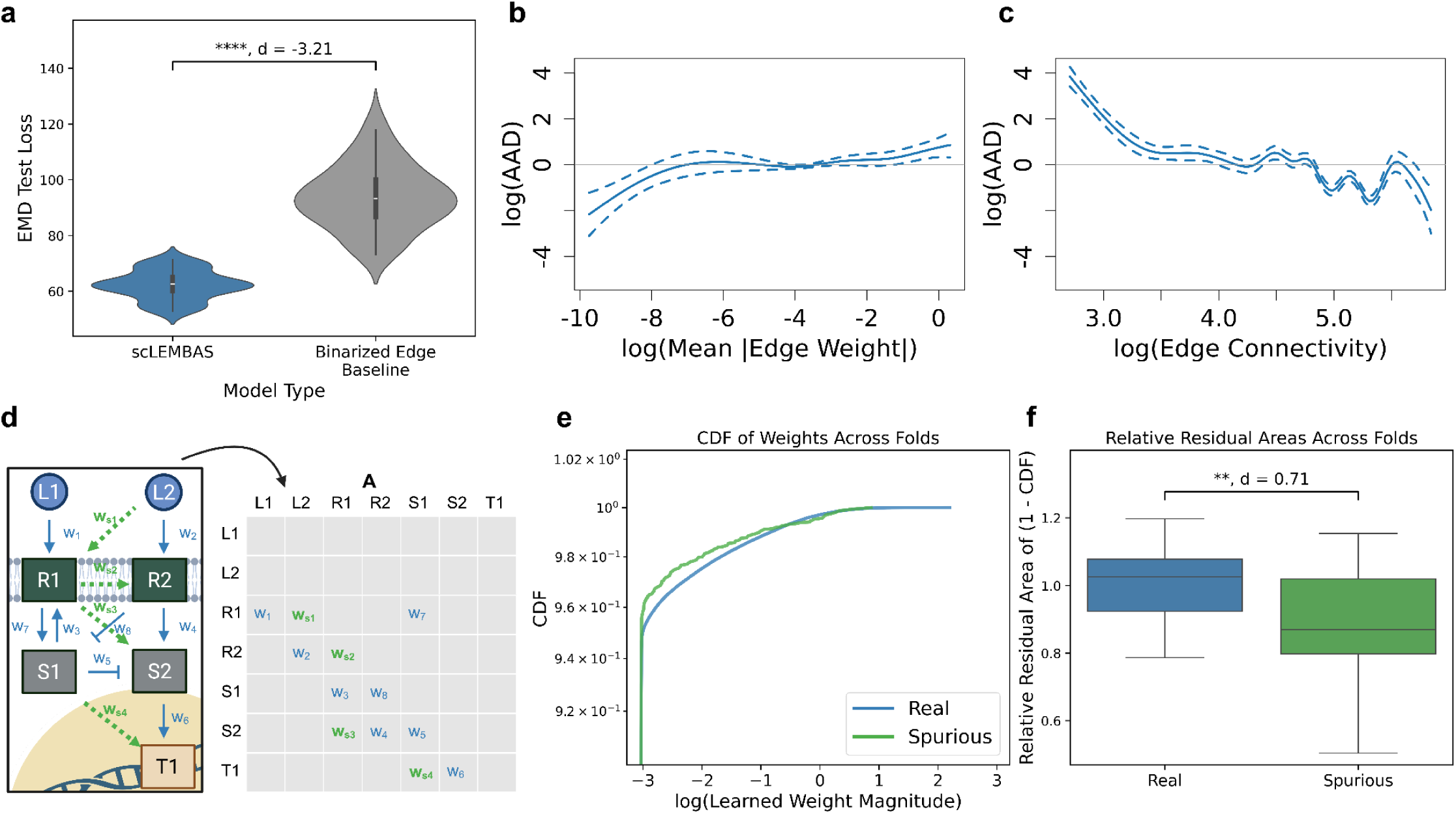
Learned adjacency matrix weight magnitudes are biologically relevant. **(a)** Violin plots compare the distribution of test EMD loss between the actual model and binarized baseline model. A one-sided paired Wilcoxon test was performed to assess whether the test loss of the actual model is lower than that of the baseline (**** p ≤ 1e-4) and the comparison is annotated by a paired Cohen’s d effect size. 10 ensemble models were generated per fold from the McCauley dataset 5-fold CV, resulting in 50 models per model type. **(b)** Fitted partial effects from the GAMM showing the association of AAD with removed edges’ mean absolute weight and **(c)** connectivity. **(d)** Graphical illustration of the added spurious edges (green) to the prior knowledge network’s “real” edges (blue), yielding additional learnable parameters in scLEMBAS’ adjacency matrix “**A**”. **(e)** CDFs are computed from the absolute values of learned edge weights separately for each of the real and spurious interactions. For visualization here, CDFs are calculated on log-transformed values across all ensembles. Only the upper quartile of edge weights is shown, representing the most biologically relevant regime as well as the region where separation is most evident. **(f)** Boxplots compare the CDF residual area between real and spurious edges. Here, CDFs are calculated separately for each ensemble model. The panel is annotated with the paired Cohen’s d and a one-sided Wilcoxon p-value (** 1e-3 ≤ p < 1e-2) testing whether the residual area of real edges is greater than that of spurious edges.

To therefore explicitly isolate the specific contribution of learned weight magnitudes beyond topology alone, we employed a linear modeling approach. Specifically, we removed a subset of edges from our learned adjacency matrix and assessed the resulting effect on scLEMBAS’ prediction. We measured the effect using the mean absolute error (MAE) comparing the baseline prediction without removed edges to that with removed edges, previously termed the “absolute average deviation” (AAD)^11^. Finally, as previously, we fit generalized additive mixed models (GAMMs) with AAD as the response, modeling the non-linear effects of learned edge weight and edge connectivity as the predictors.

We found that, after accounting for network topology, edge weight magnitude shows a significant association with AAD (F-test p = 5.35e-7). We confirmed this by quantifying the Akaike Information Criterion (AIC) of this model relative to that of a reduced model that excluded edge weight, finding a difference of 29.9.These results together indicate that learned weights capture significant and substantial additional biological information beyond network topology. Furthermore, we observe that the partial effect of weight magnitude has a positive relationship with AAD, indicating that larger weights impact model prediction more (Fig. 3b). Interestingly, however, edge connectivity showed a negative relationship with AAD, possibly reflecting the robustness of alternative paths between perturbation input and signaling activity output as a function of edges lying near high-degree nodes (Fig. 3c). Finally, the residualized number of edges (i.e., the number of edges beyond the expectation given connectivity; see Methods for details) has a strongly positive relationship with AAD (Fig. S17a). Altogether, these results demonstrate that the adjacency matrix learned weight magnitudes are relevant beyond network topology. We note that the GAMM inputs (i.e., ensembles, removed edges, and AAD) for linear model fit are the same as those used for the self-pruning analysis further described in the subsequent section.

### scLEMBAS’ Adjacency Matrix “Self-Prunes” Spurious Interactions

Given the positive association between weight magnitude and AAD, we asked whether scLEMBAS is capable of downweighting (“self-pruning”) biologically irrelevant interactions. This points towards the capability of such a model to identify signaling mechanisms involved with particular perturbation responses and biological contexts^11^ and even to identify false positives in the PKN.

To assess this, we emulated our previous approach^11^: prior to training, we introduced additional edges into the PKN that represent a baseline set of spurious interactions (Fig. 3d). Spurious edges were added in a manner that conserves the connectivity distribution of the graph, and they constituted 1% of the “real” interactions encoded by the graph. Across the 5-fold CV for the McCauley dataset, we constructed an ensemble of 5 models per fold, each initialized with a different set of spurious edges. Ensemble training largely followed the procedures used for other models trained on the McCauley dataset, but achieving self-pruning required L1 rather than L2 regularization (see Supplementary Results for details). Following training, we assessed whether spurious interactions were down-weighted compared to the real interactions. Inspection of cumulative distribution functions (CDFs) revealed a shift toward lower weights for spurious interactions in the upper quartile of weight magnitudes (Fig. 3e). We quantified this using the residual area metric previously described^11^, defined as the area between the CDF and the point (0, 1.0), such that larger value implies larger weights. Using this metric, we found that, across the entire range of weight magnitudes, the relative residual area for real edges is significantly larger than that for spurious edges, with a medium effect size (Fig. 3f).

Extending our GAMM analysis to control for edge type, we found that real edges generally had greater AAD than spurious edges—except in the highest weight-magnitude range (∼75th percentile and above), where retained spurious interactions had a substantial impact on prediction. Together with our self-pruning results in this percentile range (Fig. 3e-f), this tells us that while scLEMBAS tends to downweight spurious interactions in the high-weight magnitude range, in instances where large spurious interaction weight magnitudes are retained, they have a substantial impact on model prediction. Further details are provided in the Supplementary Results.

### Categorical Bias Identifies Nodes that Define Cell-Type-Specific Perturbation Responses

We finally evaluated the biological relevance of the categorical bias, which determines the cell type specificity of signaling pathway activity. To explore this, we selected two cell types in the McCauley dataset with distinct responses to the same perturbation: TGFB1-perturbed basal and club cells (Fig. 4d, left panel). These represented a pair with distinct perturbation shift geometries relative to control (cosine distance = 0.68), substantial separation between cell types under perturbation, and sufficient cell numbers in all four cell type - perturbation combinations.

**Figure 4:**
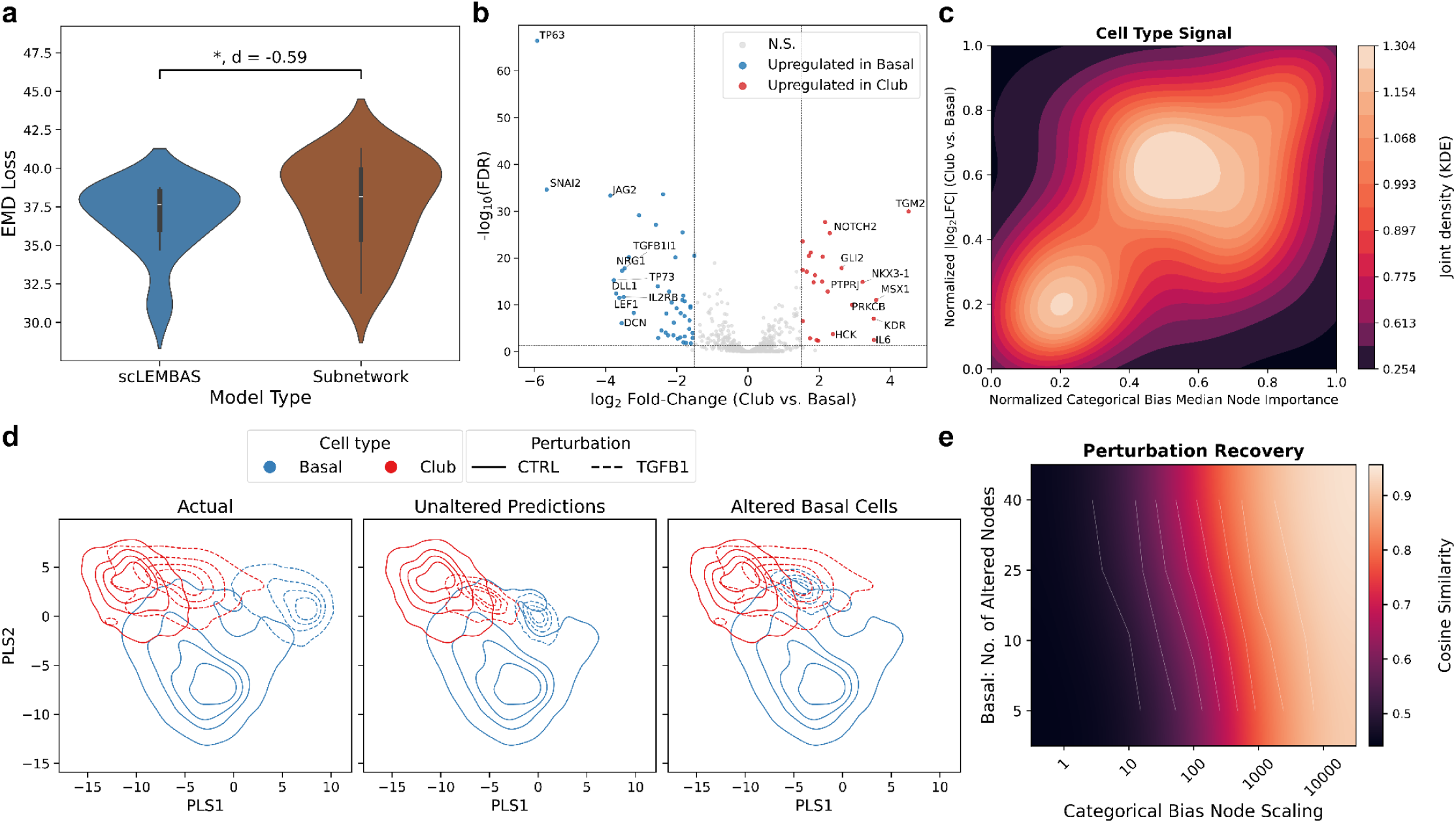
Categorical bias values are associated with cell-type specific perturbation response. **(a)** Violin plots compare the distribution of EMD loss between the actual model and the subnetwork model. Comparisons are annotated with a paired two-sided Wilcoxon signed-rank test (* 0.01 < p ≤ 0.01 ≤ 0.05) and a paired Cohen’s d effect size. 20 of 50 ensembles’ predictions, from the first and fifth fold of the original McCauley 5-fold CV, were retained for this analysis as they represented splits in which the TGFB1-perturbed club and basal cells, respectively, were in the test split. **(b)** Differential expression between club and basal cells in an external human lung dataset^25^. Volcano plot showing log₂ fold-change (LFC) between club and basal cells versus −log₁₀ adjusted p-value. Genes with adjusted p-values < 0.05 and absolute LFC > 1.5 were considered significantly differentially expressed, with genes upregulated in club cells shown in red and genes upregulated in basal cells shown in blue. The top 10 significant genes in each direction by LFC are labeled. **(c)** Two-dimensional kernel density estimate (KDE) plot illustrating the joint distribution of normalized categorical bias node importance ranks (x-axis) and normalized differential expression absolute LFC effect size (y-axis). Each axis represents per-feature ranks scaled between 0 and 1. The KDE contours depict smoothed probability density estimated over the bivariate rank space. The colorbar indicates the estimated joint probability density of observing particular combinations of normalized transcript abundance and flux ranks. Brighter colors correspond to higher densities, highlighting regions where LFC effect size and node importance co-occur more frequently. **(d)** Two-dimensional kernel density estimate (KDE) plots of club and basal cells in the first two latent variables of the PLS-reduced space. Solid lines represent unperturbed cell types, and dashed lines represent TGFB1-perturbed cell types. Unperturbed cell types always correspond to the actual data. PLS was fitted using the actual data, after which predictions were projected into the learned latent space. Plots visualize the actual data (left panel), the model predicted TGFB1 perturbed cell states (middle panel), and the model predictions upon categorical bias alteration to basal cells, with TGFB1-perturbed club cells displayed for the actual data (right panel). For categorical bias alterations, select parameters from the parameter sweep were chosen for representative visualization: the scaling factor was set to 1e4, using the top five candidate proteins. **(e)** Heatmap of cell-state recovery (defined by cosine similarity; see Methods for details) between predicted basal cells under TGFB1 with altered categorical bias node values and actual TGFB1-perturbed club cells. Alterations were conducted across a grid of perturbation parameters: the number of altered nodes (y-axis) and the node scaling factor (x-axis). Results are smoothed via bicubic interpolation and Gaussian filtering.

To assess the accuracy of the learned parameters, we generated an ensemble of models to identify a subnetwork that best defined the TGFB1 response in these two cell types (see Methods for details). This subnetwork was substantially reduced relative to the full network, containing 5,394 edges and 726 nodes, compared to 17,597 edges and 2,504 nodes in the full network. While the test loss of the subnetwork was significantly higher than that of the full network (Fig. 4a), with a moderate effect size (paired Cohen’s d = 0.58), the discrepancy in predictive performance was negligible compared to that between the full model and random baselines (Fig. 1c, EMD loss). This indicates that, despite a substantial reduction in parameters, the subnetwork still achieves reasonable predictive performance.

We next set out to determine whether we can use this to further identify specific proteins associated with cell-type specific perturbation responses. Within this subnetwork, we rank-ordered each node’s importance in the model according to the categorical bias (see Methods for details). We then compared these values to log-fold-change (LFC) effect sizes of significantly differentially expressed genes (LFC > 1.5, adjusted p-values < 0.05) between club and basal cells (Fig. 4b) from an external dataset of human lung^25^. We observed a moderate agreement between node importance and LFC effect size (Fig. 4c; Spearman correlation = 0.32), indicating that the learned categorical bias is capturing cell-type specific differences represented in transcriptomics. This is notable because signaling network node parameters are not explicitly constrained by or calibrated to expression data; thus, the model learns it simply by optimizing prediction accuracy and the integration of the various architectural components. The node importance - effect size agreement decreased when considering all significantly differentially expressed genes (any LFC value, adjusted p-values < 0.05) and all subnetwork nodes (Spearman correlation = 0.17 and 0.08, respectively), suggesting that scLEMBAS strongly emphasizes the cell-type specific signal in the most distinct nodes rather than distributing uniformly across all nodes.

Finally, using the 49 DE genes (excluding TFs and perturbations) in our subnetwork, we identified a rank-ordered list of candidate proteins to test whether their alteration, by *in silico* perturbation, caused a shift in cell state upon TGFB1 perturbation. We rank-ordered proteins by prioritizing those with high node importance while downweighting those with high centrality or absolute LFC effect size from the external DE analysis. We reasoned that this would prioritize proteins likely to cause a cell state shift without pleiotropic effects due to high centrality, and without being apparent from a simple DE analysis. Consequently, among our top five candidate proteins (MELK, MMP2, PAK5, DCN, PDGFA), only one intersected with the top 20 proteins by LFC, and only one was above the 50th percentile of centrality within the subnetwork. To implement the alteration, we replaced the learned categorical bias parameters of the predicted cell type with those of the other cell type for each candidate protein. In other words, when altering basal cells, we replaced the basal categorical bias values with the corresponding club cell values, and vice versa. Next, we conducted a parameter sweep across the top 5 to 40 candidate proteins and a range of scaling factors for the replaced values (1–1e4), assessing whether the altered cell type became more similar to the complementary cell type.

We found that this procedure successfully shifted TGFB1-perturbed basal cell predictions toward TGFB1-perturbed club cells (Fig. 4d, Fig. S19a). In contrast, we were not able to strongly alter club cell states (Fig. S19). This may reflect differences in cell plasticity, with basal cells serving as a multipotent progenitor population, whereas club cells represent a more downstream, fate-committed cell type. The shift in TGFB1-perturbed basal cell state was more strongly driven by the scaling factor than by the number of altered nodes (Fig. 4f, Fig. S19a), indicating that a minimal set of strongly altered proteins can drive cell-type- and perturbation-specific alterations to cell state. Notably, altering the categorical bias values did not make the basal expression shift geometry from control to perturbed consistent with (i.e., parallel to) that of club cells, but instead altered the final perturbed cell state.

Altogether, these results indicate that the categorical bias captures biologically meaningful cell-type-specific signaling differences and can be used to prioritize proteins whose alteration drives cell-state shifts.

## Discussion

Here we present scLEMBAS, a gray-box neural network approach that extends the mechanistic LEMBAS framework^10^ to model signaling pathway activity across biological contexts at single-cell resolution. scLEMBAS decomposes LEMBAS bias, which conceptually represents protein-specific activity, into a categorical term that encodes context—here, cell type—and a global term that captures the residual cell-to-cell heterogeneity; by adversarially removing perturbation and context information from the global bias, the model can substitute these covariates at prediction time and thereby answer, at single-cell resolution, the counterfactual of what a given cell’s transcription factor (TF) activity would have been had it received a different perturbation. Across two single-cell RNA-sequencing datasets spanning binary and multi-perturbation settings, scLEMBAS accurately predicted out-of-distribution combinations of cell type and perturbation, capturing both systemic perturbation effects and perturbation-specific directional structure as assessed by a diverse panel of complementary metrics. Targeted ablations confirmed that each architectural component—the adjacency matrix, the categorical bias, and the global bias—contributes meaningfully and in its intended conceptual role. Notably, the global bias not only enabled accurate counterfactual separation of perturbations from their controls but also retained subtype-level variance that supported subtype-specific perturbation responses.

A central advantage of scLEMBAS is that its predictions are grounded in interpretable, mechanistically meaningful parameters rather than opaque latent representations. Because perturbation effects propagate through a prior-knowledge signaling network whose edge weights are explicitly learned, the model’s parameters can be interrogated directly. We found that the learned adjacency weights carry biological information beyond network topology alone: weight magnitude was significantly associated with predictive influence, and scLEMBAS preferentially down-weighted (*“self-pruned”*) spurious interactions relative to curated ones, suggesting a capacity to identify false positives in the PKN and to nominate the interactions most relevant to a given perturbation response. Similarly, the categorical bias captured cell-type-specific signaling differences that agreed with independent transcriptomic measurements and could be altered *in silico* to prioritize a compact set of proteins whose perturbation drove cell-state shifts. Together, these results establish scLEMBAS as a mechanistic, genome-scale, and context-aware model of single-cell signaling activity.

In comparisons against pseudobulk baselines, we observed that both accounting for cell type in addition to the perturbation effect—the original goal of scLEMBAS and similar models—as well as evaluating within a reduced latent space that appropriately defines the variance of interest improved scLEMBAS’ relative performance. This highlights the importance of selecting relevant assessments of model performance. Broadly, single-cell RNA-sequencing captures global measurements of mRNA abundance, whereas perturbation effects may be sparse, particularly relative to cell-type-driven variance^7,26^. Consequently, it is crucial to both define the expected changes (e.g., in a data-driven manner via reduced latent spaces) and how they should be quantified (e.g., global metrics across all features may not be as informative for targeted, highly-specific perturbations with local effects). Accounting for cell type underscores the limitations of assessing single-cell perturbation models using perturbation means alone: depending on the model’s predictive task, ignoring cell-type context can reduce predictive resolution and distort performance comparisons, and future baseline models should therefore incorporate context when relevant to enable more appropriate and informative evaluations, despite their goal of being “deliberately simple”. The improved performance observed in the reduced space further emphasizes the importance of evaluation metrics for single-cell perturbation modeling, an area in which the field is still actively converging^16,17,27,28^. Reduced representations preferentially capture biologically relevant signals, whereas standard, global machine-learning metrics applied in the full feature space are largely agnostic to this and may consequently obscure such signals. Together, our findings suggest that both model design and evaluation strategy critically shape conclusions about relative model performance, and it is when scLEMBAS’ additional capabilities—accounting for network topology, learning the weights of the interactions, and capturing cell-type dispersion—are taken into account that its utility over simple baselines fully emerges.

Several limitations temper these conclusions and motivate future work. First, although adversarial training substantially reduced cell-type and perturbation information in the global bias, removal was incomplete and sensitive to hyperparameters. Similarly, fully disentangling categorical-context from perturbation information remains an open challenge^7^; the shared dimensionality between the bias and the signaling network may be unnecessarily high, and initializing the categorical term with pre-trained cell-type embeddings could impose a more informative, lower-dimensional prior. Second, achieving self-pruning requires L1 rather than L2 regularization; going forward, this choice may be preferable for mechanistic insight, though merits further investigation regarding predictive performance and generalization across datasets. Third, we were unable to recover edge directionality above chance when blinding the model to directionality in a small fraction of edges, suggesting that our current perturbation coverage does not provide sufficient signal to resolve interaction directionality; broader perturbation coverage across the network, potentially augmented by incorporation of endogenous cell-cell communication^29,30^ signals, may help recover this signal. Finally, our analyses of the multi-perturbation McCauley dataset required excluding perturbations with negligible transcriptomic effect (see Methods and Supplementary Information for details); the limited effect of some perturbations on global single-cell measurements has been observed previously and is known to reduce the predictive performance of single-cell foundation models^26,31,32^. Ideally, such perturbations would be handled within training itself—by making the model aware that some perturbations may exert little or no effect and accounting for this accordingly—rather than removed *a priori*.

Beyond addressing these caveats, scLEMBAS’ mechanistic architecture opens several natural extensions. Because perturbations are encoded through the signaling network via learned parameters rather than as a latent embedding, scLEMBAS may support zero-shot perturbation prediction—generalizing to conditions in which the perturbation itself, and not only its pairing with a context, is out-of-distribution. This would require the corresponding input perturbation edge to be specified in the network. This capability is unavailable to purely embedding-based models. The same mechanistic encoding naturally accommodates intracellular perturbations which we did not explore here but are of particular relevance to therapeutic target discovery. Relatedly, the perturbation input as currently formulated encodes only the presence or absence of a single perturbation at a time; combinatorial perturbations and continuous, dose-dependent values are not yet supported, but both could be accommodated by modifying the discriminator architecture. As mentioned before, incorporating endogenous cell-cell communication would further extend the model beyond exogenous perturbations to the signaling milieu that cells generate for one another, enriching the information from which the model can learn the adjacency matrix weights from and the biological realism of predicted responses.

Overall, scLEMBAS simultaneously provides genome-scale, mechanistic simulations across multiple contexts at single-cell resolution. By coupling the interpretability of prior-knowledge signaling networks with the resolution of single-cell measurements and the counterfactual flexibility of a context-decomposed neural network, scLEMBAS enables quantitative dissection of how signaling pathway activity is reshaped by perturbation within specific cellular contexts. This is especially pertinent to settings in which the heterogeneity of cellular responses governs outcomes, where identifying the specific pathway components can enable tuning of cell state toward favorable phenotypes.

## Methods

### Single-Cell RNA-Sequencing Data Preprocessing

We downloaded raw counts for two single-cell RNA-sequencing datasets: the “Kang” dataset^14^ and the “McCauley” dataset^15^ from Seurat and Zenodo (GSE246368; https://zenodo.org/records/10602177), respectively.

Preprocessing was conducted with scanpy^33^ with default arguments unless otherwise specified.

The Kang dataset was filtered for cells containing fewer than 100 genes, and genes detected in fewer than 3 cells were removed from the dataset. Erythrocytes, plasmacytoid dendritic cells, and megakaryocytes were excluded from downstream analysis, representing fewer than 2% of the cell population (≤ 236 cells). Counts were normalized for depth and scaled by 1e6 to attain counts per million (CPM). We add a pseudo-count of 1 to the CPM values and log-transformed them. The top 5000 highly variable genes (HVGs) were retained using the dispersion-based method as described here^34^ (‘flavor = ‘seurat’’).

The McCauley dataset was filtered for cells containing fewer than 6 genes, and genes detected in fewer than 3 cells were removed from the dataset, recapitulating the original thresholds^15^. PNECS, tuft cells, and ionocytes were excluded from downstream analysis, representing fewer than 1% of the cell population (≤ 242 cells); cells that were not assigned a cell type were also removed. Of the 68 remaining cell type - perturbation combinations, 7 contained fewer than 100 cells and were also excluded from downstream analysis. Normalization and HVG selection was conducted the same as in the Kang dataset.

Transcription factor (TF) activity for each dataset was inferred from log-normalized HVGs using decoupler^13^. The CollecTRI^35^ gene-regulatory network was used, with TFs containing fewer than five target genes in the dataset excluded. The final activity score was taken as the consensus across the multivariate linear model, univariate linear model, and weighted sum, as described by decoupler^13^.

For the McCauley dataset, upon TF activity inference, we noticed that perturbations had the weakest covariate signal of data-driven variance relative to cell type, cell cycle, and batch donor. (Fig. S3). Training on perturbation labels that have sparse effects on transcriptomics relative to perturbation-invariant information allows the model to achieve reconstruction while disregarding perturbation information. Consequently, we subsetted to the five perturbations that we detected to have a significant separation from control, unperturbed cells (TGFB1, IFNG, IFNA2, BMP4, IL13). For an extended discussion of methods and reasoning on this subsetting, see the Supplementary Information.

### Signaling Network Reconstruction

Physical, human protein-protein interactions were retrieved from Omnipath^9^ (retrieved May 24, 2024). The McCauley dataset contained a couple perturbations that were not present in the dataset, and these were added according to a manual literature search: an inhibiting edge was added between the CHIR99021 perturbation and each of GSK3A, GSK3B, CDK1, and MAPK targets; activating edges were added between the TNF perturbation and TNFRSF1A and TNFRSF1B targets. The network was filtered to exclude interactions with a curation effort score less than two and fewer than one associated reference. Self-loops within a single node were also excluded. Next, the network was filtered for those interactions that provide full paths between perturbations and transcription factors. Specifically, all nodes that connected both to sources (perturbations) and targets (TFs) were retained. The final signaling network for the McCauley dataset contained 17,597 interactions between 2,504 proteins, retaining 380 of 423 TFs. The final signaling network for the Kang dataset contained 17,548 interactions between 2,478 proteins, retaining 336 of 418 TFs.

### Latent Space Embeddings

To assist in visualization and quantitative assessment of model predictions, we embedded TF activity in linear latent spaces. We applied principal component analysis (PCA) to the Kang dataset. For the McCauley dataset, we applied Partial Least Squares (PLS) due to the aforementioned reduced signal in the covariates of interest. Predictions were always projected into the latent space fit on the actual data, rather than being jointly embedded.

When applying PCA, unless otherwise stated, all analyses were conducted using scanpy with default arguments. The number of principal components was automatically selected using the Python package *kneed*, which identifies the elbow of the cumulative variance versus number of components curve. The neighborhood graph was constructed in PCA space with 15 neighbors, and downstream UMAP was fitted on the PCA space. For Leiden clustering, the resolution parameter was selected to maximize the normalized mutual information between the cell type-perturbation combination label and the clusters, emulating previous approaches^36^. Resolutions between 0.01 and 3 were tested.

When applying PLS, we fit *sklearn* PLS models to specified one-hot encoded discrete responses (cell type, perturbation, or cell type - perturbation combinations) depending on the specific question of interest. This enabled visualization, as well as quantification, in instances where standard PCA did not provide the desired separation by categorical label. Regardless of the response variable, we used the same pipeline to fit PLS models. First, we identified the appropriate number of PLS components. To do so, we fit a model at each component between 1 and 25 components and assessed the mean 5-fold cross-validation (CV) classification accuracy. Next, using the Python package *kneed*, we automatically identified the elbow of the accuracy vs. number of components curve. Finally, we ensured the model fit was statistically significant by checking that the 5-fold Q^2^Y is greater than (one-sided *p* ≤ 0.05) a distribution of 100 null Q^2^Y values obtained from models fit on permuting the response labels. PLS were fit on specific subsets of data depending on the evaluation of interest, as elaborated on in the Supplementary Methods.

### Model Structure and Prediction

#### Structure

The predictive objective of scLEMBAS is to estimate transcription factor (TF) activity for out-of-distribution (OOD) “conditions”: unseen combinations of cell type (context) and perturbation, in which each individual cell type and each individual perturbation was observed during training but their specific combination was not. Following the compositional logic established for the bulk LEMBAS framework and the fader-network–inspired decomposition of CPA, scLEMBAS answers a counterfactual question at the single-cell level — what would the TF activity of a given cell have been, had it been exposed to a different perturbation than the one it actually received — by recombining the learned components of the model under the target condition.

This is made possible by the architecture of scLEMBAS, which is modified from the described GPU-adaptation of LEMBAS^11^ to account for the decomposition of bias into global and categorical components. The context-independent protein–protein interaction strengths are encoded in the signaling-network weights (the adjacency matrix), while protein-specific activity is captured by the additive bias (Fig. 1A). Because the global bias is produced by a stochastic encoder from a cell’s gene-expression vector with perturbation and context information adversarially removed during training, it carries the residual, cell-to-cell heterogeneity that is independent of both the perturbation and the cell type. The perturbation and context are therefore free to be substituted at prediction time, leaving the cell’s individual basal state intact (Fig. S1). The conceptual grounding of each architecture component is discussed extensively in the Results section. All model construction and analysis was conducted using *PyTorch*^37^.

#### Forward pass

The target perturbation input was passed through the input projection layer, which scales the perturbation input by the input projection amplitude and places it into the full set of signaling-network nodes, yielding the perturbation contribution to the network.

The categorical bias is supplied by a learnable embedding for each categorical covariate; given a covariate label, the corresponding embedding indexed by the covariate value is added to the node-wise bias. As with all bias terms, the entries corresponding to perturbation input nodes are masked to zero so that bias remains independent of the perturbation input. The global bias was obtained by passing the source cell’s gene-expression vector through a Gaussian variational encoder, comprising fully connected hidden layers that output a per-node mean and log-variance, from which the global bias is drawn via the reparameterization trick^38^, with the standard deviation floored by a small constant for numerical stability.

The total bias was formed as the sum of the global bias and the categorical bias, and this total bias was added to the projected ligand input to give the combined input-plus-bias term that drives the recurrent network. The signaling state was then propagated to steady state with the recurrent formulation of LEMBAS: the hidden state was initialized at zero and updated iteratively by multiplying the current state by the signaling-network adjacency weight matrix, adding the combined input-plus-bias term, and applying the Michaelis–Menten–like (MML) activation function. The activation constrains node states to a physiologically interpretable range and, through the recurrent structure, allows the network to capture feedback loops. Iteration proceeded until the maximum change in node state between successive checkpoints fell below a convergence tolerance, indicating that a steady state had been reached.

The steady-state activity over all network nodes was passed through the output projection layer, which selects the TF nodes and applies a learned linear scaling and bias, to produce the predicted TF-activity vector.

#### Constructing counterfactual inputs

For each OOD test condition, we identified a corresponding source condition in the training data from which to ask the counterfactual. In most instances here, the source shares the same cell type as the test condition but carries the control, unperturbed cells; for the Kang dataset, in which the perturbation is binary, we tested forward passes in both directions: from control to perturbed and vice-versa. Predicting unperturbed cells effectively tests the contribution of bias terms independent of the adjacency matrix. For every cell belonging to the source condition, three model inputs were assembled. First, the cell’s gene-expression vector was used as the input to the stochastic encoder that generates the global bias. Second, the perturbation input was set to the target perturbation of the test condition, encoded as a one-hot dummy vector. Third, the categorical covariate index encoding the cell type was set to that of the test condition, retrieving the corresponding embedding. Inputs were accumulated across all cells in the source condition.

#### Train-test splits

Train-test splits were strategically generated such that OOD test conditions were unseen during training, but the individual cell types and perturbations were seen (no zero-shot test conditions). 5-fold splits were performed at the level of conditions, tolerating a 10 and 5 percentage point population frequency deviation (for the Kang and McCauley datasets, respectively) in the 80%-20% train-test split at the level of individual cells to meet the remaining criteria. For the McCauley dataset, control, unperturbed cells were excluded from the splitting procedure to ensure they are all present in training. To ensure sufficient coverage of perturbation and cell type, we ensured 50% and 60% of all conditions containing the given perturbation and cell type, respectively, were present in training. For the Kang dataset, we binned cell type labels by their counts into equal-size quartiles and ensured each test split contained atleast one cell type from atleast three of the four bins.

### Model Training

For each cell, the forward pass is run without a counterfactual – the cell under the same cell type and perturbation is predicted, and projected to transcription factor (TF) activities, which are compared against the inferred TF activities. As in the GPU implementation, the model is trained with the Adam optimizer using a warmup phase followed by cosine annealing with warm restarts, the signaling weights are masked to the PKN topology after every update so that only edges supported by the PKN remain non-zero, Gaussian noise is injected into the signaling-network input and into the parameter gradients to improve robustness, and the edge sign (activating or inhibiting) is softly constrained by an L1 penalty on weights whose sign conflicts with the PKN annotation. The signaling weights, the output projection weights, and the output projection bias were additionally regularized with L2 penalties. Unlike the original formulations, the spectral-radius and uniform state regularizations were found to be unnecessary for stable training in the GPU implementation and were therefore disabled. Next, we describe the components that are specific to scLEMBAS.

#### Categorical and Global Biases

The latent global bias is regularized toward a standard normal prior with a scaled KL-divergence term. The encoder’s linear layers are separately regularized with a small L2 penalty. The encoder parameters are excluded from the optimizer used for the remaining model parameters and are instead updated by a dedicated Adam optimizer with its own learning-rate schedule; this is purely a separation of optimizer state and learning rate, not of objective, since both optimizers act on gradients from the same loss.

#### Regularization between categorical bias and perturbation

Because the categorical bias and the perturbation input are both additively combined within the signaling network, the categorical bias can in principle absorb perturbation-specific signal, which would undermine the intended factorization of context and perturbation. The categorical embeddings are constrained to a maximum L2 norm and are regularized with an L2 penalty, which discourages the categorical bias from collapsing onto memorizing perturbation-specific structure and instead encourages generalization across context–perturbation combinations. To further explicitly discourage this, we add an orthogonality regularization between the categorical bias terms and the perturbation input. The penalty captures linear dependence between the two via a Frobenius-norm-based criterion and is added to the model objective, complementing the L2 penalty on the categorical embeddings.

#### Adversarial disentanglement of the global bias

To make the global bias counterfactually exchangeable across conditions, context (cell type) and perturbation information are adversarially removed from it, following the fader-network strategy used in the compositional perturbation autoencoder (CPA)^6^. Two discriminators are trained to predict, from the global bias alone, (i) the categorical covariate label and (ii) the perturbation label. The discriminators are multilayer fully connected classifiers with spectral normalization, dropout, and label smoothing. Each discriminator is trained with cross-entropy (or binary cross-entropy for two-class problems) against the detached global bias for several inner steps per generator update, with gradient-norm clipping.

The encoder is trained adversarially against these discriminators so that the global bias becomes uninformative of context and perturbation. Rather than maximizing the discriminator loss directly, we use the label-flipping variant: the adversarial loss is computed against flipped labels, which are sampled in proportion to the empirical training class frequencies. The two adversarial losses are added to the model objective with a per-epoch penalty weight that is ramped up over training following a power curve, so that the disentanglement pressure increases gradually as the model learns. To allow the model to first learn the perturbation and categorical structure before the generator and gene-expression encoder are engaged, adversarial training begins only after a fixed number of warmup epochs.

#### Loss and optimization

The full training objective for the model (excluding the separately optimized discriminators) is the sum of the reconstruction loss and the regularization terms described above. The reconstruction loss is the mean squared error between predicted and inferred TF activities, with no counterfactual. To prevent abundant conditions from dominating the gradient and to give rarer conditions proportionate influence, the reconstruction loss is computed separately for each unique context–perturbation condition within a batch and combined as a weighted average, with weights proportional to the square root of the number of cells per condition. The reconstruction loss is scaled by a constant so that it remains on a comparable magnitude to the regularization and adversarial terms.

Each minibatch update therefore proceeds in two stages. First, the categorical and perturbation discriminators are updated on the detached global bias. The remaining model is then updated on the combined reconstruction, regularization, KL-divergence, orthogonality, and adversarial losses: this combined loss is backpropagated once, and the resulting gradients are applied by two optimizers in the same step — one for the global-bias encoder and one for all other trainable parameters (signaling weights, categorical embeddings, and output projection), each with its own learning rate. Gradient noise is injected and the masks are re-applied after each update.

Details of specific training dynamics across epochs are described in the Supplementary Results. Specific hyperparameters are delineated in Table 2. Hyperparameters were selected empirically and tuned for convergence during training because the models’ training is sensitive to adversarial information removal; multiple training objectives, including minimization of predicting errors, a warmup period to allow the discriminators to learn labels, prevention of mode collapse by the generator, adversarial information removal by the generator, and appropriate other regularizations (e.g., of edge weight directionality) simultaneously must be accounted for.

### Baseline Models

At single-cell resolution, we generated three baseline models with the same architecture and hyperparameters as scLEMBAS. The first was a “random” baseline trained on permuted transcription factor features. The second was a “no adversarial removal” baseline, in which the adversarial penalties for both cell type and perturbation were set to zero throughout training, removing any explicit pressure on the generator to eliminate information about these categories.

The third was a binarized baseline for topological assessment. For each split in the McCauley dataset 5-fold CV, we created an ensemble of 10 models. Each ensemble was trained with different seeds, yielding differences in parameter initialization values as well as any additional stochastic components of the training (e.g., generator reparameterization trick^38^); all other components of the model and training dynamics were held constant. This ensemble of 50 models is also used in the subnetwork analysis subsequently described. For each trained model, we generated a corresponding baseline model with binarized edge weights, representing a topology-based network indicating simply whether an interaction exists between two proteins or not. The value of the edge weight for interactions that exist was set to the mean of the absolute learned values of the corresponding actual model. To make comparisons more stringent, we also retained interaction directionality (activating or inhibiting): in the case of known directionality from the PKN, we ensured signs were correct (positive for activating, negative for inhibiting); in the case that directionality was not specified from the PKN, we ensured the directionality matched that learned by the actual model. EMD test loss was calculated as previously described, taking the mean across each test condition for a given split.

### Evaluation Metrics to Assess Model Predictions

Cosine distances were computed following the metric described by Viñas Torné et al.^18^. Briefly, this metric quantifies the angular distance between two vectors defined relative to a shared control reference. The control vector is defined from the control centroid to the mean centroid across all perturbations, capturing global, non-specific perturbation effects. The perturbation-specific vector is defined from the same control centroid to the centroid of an individual perturbation. Larger cosine distances indicate greater perturbation-specific deviation relative to shared, systemic effects.In our analysis, this computation was performed separately for each fold and restricted to held-out test conditions and their corresponding controls. Accordingly, both the set of perturbations and the control cells used to define centroids were limited to those present in a given test split.

The Earth Mover’s Distance (EMD) was calculated using the ‘SamplesLoss’ method from the Python package *geomloss*, initialized with default parameters. EMD loss was calculated for each condition separately, and averaged across conditions. For OOD prediction accuracy, the mean across test conditions was taken for each fold. For ablation experiments, scLEMBAS was trained with all components, and each ablated component was excluded during the forward pass and compared to the model with all components.

The rank-based metric was implemented as described in Wu *et al*.^17^, with the final score subtracted from one so that larger values indicate better predictive performance, with 1 being a perfect score. Briefly, predicted and observed perturbations were first pseudo-bulked by averaging expression. Next, distances between all predicted–observed perturbation pairs were computed within each test condition using the Manhattan distance as in the Virtual Cell Challenge^16^. Finally, each predicted perturbation was ranked by its relative distance to the correct corresponding observed perturbation relative to all other predicted perturbations. The mean across all predictions was taken to get the final score. For OOD prediction accuracy, within each fold, the rank score was computed as the mean across test perturbations

### Differential Feature Concordance

Significantly differential features were identified using a Mann–Whitney U test comparing predicted perturbed cells to their respective controls from the actual data for each test condition, with p-values adjusted for multiple testing using the Benjamini–Hochberg false discovery rate (FDR) correction. Features with a FDR ≤ 0.05 were retained as significantly differentially expressed. Significantly differentially expressed were rank-ordered by Cliff’s delta effect size and separated by those with positive (upregulated) or negative (downregulated) effect sizes. This was repeated for each test condition across all 5 folds, comparing both predicted and actual data perturbations to the respective actual data control. To control for differences in statistical power arising from unequal numbers of cells in predicted versus true perturbations, we subsampled both to the minimum shared cell count per condition. This procedure was repeated 100 times to obtain robust estimates of concordance.

Next, agreement between predicted and actual differential expression results was assessed using a modified concordance at the top analysis^22,23^. At each feature rank, we computed the Jaccard index between the sets of predicted and true differentially expressed features, considering all features at that rank or higher. Each comparison (predicted or actual) is limited by that with fewer significantly expressed features.

To synthesize concordance across test conditions and effect size directions, we used a generalized additive mixed model (GAMM). The GAMM was fitted using the R package *mgcv* (v1.9-4) with the number of basis functions for smooth terms set to *k* = 5. We modeled concordance as the response, and included a fixed effect for the model type (scLEMBAS and the baselines). To allow concordance–rank relationships to vary by model, we fit model type specific smooths over rank via an interaction, yielding separate rank-dependent curves for each model type. We also included a fixed effect for the effect size direction, and random effects for the test condition and subsample iteration. Fitting was restricted to the top 50 ranked features to ensure comparisons focused on consistent, biologically meaningful signals. Partial effects and F-test p-values were obtained from models fit using fast restricted maximum likelihood (fREML), enabling stable estimation of smooth and random effects in large datasets.

To directly quantify differences in concordance between scLEMBAS and baseline models across the rank spectrum, we computed model-based contrasts using the fitted GAMM. For each rank, we estimated the expected difference in concordance between two model types as the contrast of their fitted linear predictors, marginalized over effect-size direction. Formally, this contrast was defined as the expected difference in fitted concordance between two model levels, averaged across effect-size direction in proportion to the observed frequency of each direction in the data. Contrasts were computed between model-specific fitted concordance curves rather than individual coefficients. Specifically, we constructed contrasts at the level of the linear predictor, allowing differences between models to be evaluated across the full smooth function of differential expression rank. Uncertainty was propagated using the full coefficient covariance matrix, yielding rank-wise Wald confidence intervals that jointly account for both parametric and smooth components shared across model types. Random-effect smooths were excluded to obtain population-level comparisons. The resulting contrast curves therefore represent the estimated difference in fitted concordance between models as a continuous function of differential expression rank. Global performance differences across feature ranks were summarized using a mean normalized area-under-the-curve (nAUC) metric. Specifically, the fitted rank-wise contrast between models was numerically integrated using the trapezoidal rule and normalized by the rank range, yielding the average fitted concordance difference across ranks.

Partial effect curves represent the model-inferred relationship between concordance and differential expression rank for each model type. For visualization, predictions were generated across ranks while holding all other covariates at fixed reference values (first factor level or median for numeric variables). As in the contrast analysis, fitted values were marginalized over effect-size direction, and random-effect smooths were excluded to reflect population-level trends.

### Cell Subtype Analysis

Club, goblet, and multiciliated cells in the McCauley dataset were further annotated by cell state as either “transitional” or “mature”.

In assessing the presence of cell subtype information in cell types, we first obtained the global bias output for all OOD test conditions in each fold. We ran our previously described PCA pipeline on the global bias generated for a given cell type. Next, we assessed the extent to which cell subtype information was retained in this PC space by calculating the mean test-split balanced accuracy from 5-fold cross-validation using logistic regression and random forest classifiers (see Supplementary Methods - Latent Space Covariate Signal Quantification for details on the classifiers). This was run on each cell type within a test split separately in order to avoid confounding due to dependencies between the cell type and cell subtype labels.

Counterfactual predictions were run for a given cell type within a test OOD condition, stratified by their cell subtype. For example, if a test split included multiciliated cells under BMP4 stimulation, we would run the standard forward pass from the “origin” condition seen in training (e.g., multiciliated cells under no perturbation); thus the “origin” here is the unperturbed multiciliated cells and the “target” is the BMP4-stimulated multiciliated cells, which can further be stratified into “transitional” and “mature” subtypes. The subtype label of the input cells of a particular cell type in the origin was used to stratify predictions of the target. Counterfactual predictions were restricted to cases with at least 25 cells in each of the four perturbations–subtype combinations, completely excluding goblet cells. We then filtered for those cell types with at least 100 cells per subtype, excluding goblet cells from this analysis.

To visually characterize subtype expression shift geometries under perturbation (Fig. S16), PLS models were fit separately on each OOD test condition containing the cell type and perturbation pair of interest using the standard embedding pipeline described above, with one exception: perturbation and cell subtype responses were not aggregated into a single “condition,” but instead retained as separate variables. Because this was implemented via one-hot encoding rather than PLS2, dummy matrices were concatenated across the multivariate responses. Because counterfactuals predicting perturbed cells from control rarely exhibited divergent subtype shifts (see Supplementary Results for details regarding expression shift geometry), for this analysis, we also included counterfactual transitions between perturbations (from one non-control perturbation in the training split as the “origin” to a perturbation in the test split as the “target” within a cell type) across the five folds. Inclusion criteria required that both Euclidean and Mahalanobis distances between subtypes were larger in the target perturbation than in the origin perturbation (ensuring divergent shifts).

### Self-Pruning and Linear Modeling of Model Sensitivity to Edge Weights

Self-pruning was assessed using the same approach previously implemented for bulk LEMBAS^11^. Briefly, we trained an ensemble of 25 models on the McCauley dataset, comprising 5 models per fold across the previously described 5-fold cross-validation split. For each model, we introduced 176 spurious edges to the PKN prior to training, corresponding to 1% of existing interactions. To reflect the network topology and avoid introducing biases due to node degree, spurious edges were added in a manner that maintained the connectivity distribution of the PKN, following the approach described in Nordenstorm et al.^11^. Each ensemble was also trained with different seeds, yielding differences in parameter initialization values as well as any additional stochastic components of the training.

We were interested in understanding the functional relevance of various network properties, as assessed by their effect on scLEMBAS’ predictions. To this end, we used the previously described “absolute average deviation” (AAD) metric ^11^, which quantifies the change in model prediction from a baseline upon perturbation as the MAE between the two predictions. For each ensemble model generated in the self-pruning analysis, we implemented an iterative procedure. We first quantified the connectivity of all edges in the network, defined as the average degree of the two nodes connected by a given edge. We then iterated across equal-interval bins (spanning connectivity values of 15 per bin), removing all spurious edges that lie within a given connectivity bin. With these removed edges, we re-ran the model prediction and quantified the AAD as the mean absolute error (MAE) between this prediction and the model without removed edges serving as the baseline. We then removed a corresponding number of true edges within the same connectivity bin by random sampling, repeating this procedure 10 times, and recorded the resulting AADs. For downstream analysis, in each iteration, we stored the AAD alongside the connectivity bin, mean removed edge weight, number of edges removed, and removed edge type (spurious vs. true).

All generalized additive mixed models (GAMMs) were fitted using the R package *mgcv* (v1.9-4) with the number of basis functions for smooth terms set to *k* = 15. Partial effects and F-test p-values were obtained from models fit using restricted maximum likelihood (REML), while likelihood ratio tests and Akaike Information Criterion (AIC) comparisons were performed using maximum likelihood (ML) to ensure valid comparison of nested models differing in smooth terms. To model prediction sensitivity (i.e., AAD), we fit a GAMM with AAD as the response, including smooth terms for learned mean absolute edge weight, edge connectivity, and the residualized number of edges removed, and fixed effects for edge type (spurious vs real), and a random intercept for model identity. Prior to model fitting, several covariates were log-transformed: AAD, connectivity, number of removed edges, and mean edge weight. Because the number of removed edges and connectivity exhibited high concurvity, the log-transformed number of removed edges was residualized against log-transformed connectivity via ordinary least squares regression. Due to structural dependence between connectivity and the number of edges removed arising from the degree-bin removal procedure, their relative contributions are not fully identifiable, even after residualization. We therefore focus on the independent contribution of edge weight, quantified as the change in AIC upon its removal from the full model, rather than directly comparing weight and connectivity importance. To assess whether the relationship between edge weight and AAD differed by edge type, we fit a second GAMM allowing the smooth effect of learned edge weight to vary by edge type, while retaining the same covariates and random-effects structure as the main model.

### Categorical Bias Node Subset Identification and Alteration

A brief overview of our categorical bias analysis follows; full methodological details are provided in the Supplementary Methods. To assess the relevance of the categorical bias, we focused on club and basal cells in the McCauley dataset under TGFB1 perturbation, a cell-type pair with sufficient cell numbers (>75 per condition) and distinct perturbation responses (Fig. 4d). We first identified a subnetwork of protein-protein interactions defining the club and basal TGFB1 response, retaining the top 5 TFs rank-ordered by a PLS fit on a multivariate response of cell type and perturbation. Nodes were then rank-ordered by a node importance score adapting the approach of Meimetis et al.^12^, combining the absolute magnitude of the learned bias term with a measure of how the bias shifts the RNN output between control and perturbed forward passes, aggregated as the median rank across the model ensemble. To identify candidate perturbation targets, we compared node importance against differential expression effect sizes (|LFC|) computed on an orthogonal dataset (HLCA v1.0)^25^ via pseudobulk PyDESeq2 analysis contrasting club and basal cells. Focusing on significant genes that were neither TFs nor ligand perturbations, we ranked candidates by a composite priority score combining node importance (60%), inverted |LFC| (20%, to prioritize candidates missed by differential expression), and inverted network centrality (20%, to downweight likely pleiotropic nodes). Finally, we tested whether altering the categorical bias of top candidate nodes—replacing one cell type’s categorical bias values with a scaled version (1x–1e4x) of the others and re-running forward passes—could "recover" the target cell type’s TGFB1 perturbation state, quantifying recovery with cosine- and Euclidean-based similarity metrics relative to the observed data.

## Code and Data Availability

scLEMBAS package and analysis files can be found at https://github.com/hmbaghdassarian/scLEMBAS. All data was downloaded from publicly available resources: the “Kang” dataset ^14^ and the “McCauley” dataset^15^ were downloaded from Seurat and Zenodo (GSE246368; https://zenodo.org/records/10602177), respectively. The Human Lung Cell Atlas (HLCA v1.0)^25^ data was downloaded from the following link: https://data.humancellatlas.org/hca-bio-networks/lung/atlases/lung-v1-0. Initial Omnipath files used for generating the signaling networks can be found at https://doi.org/10.5281/zenodo.10823115.

## Authors, Contributions, and Acknowledgements

H.M.B. and D.A.L. conceived the work. All authors provided important insights for formulating and analyzing the scLEMBAS model. H.M.B. implemented the Python package and performed analyses of predictive accuracy and architectural component utility. N.M. generated the subnetwork. O.N. assisted with the self-pruning analysis. A.N., O.N., and N.M. provided specific algorithmic suggestions for model implementation. B.A.J. conceived of the subtype analysis. H.M.B. wrote the paper and all authors carefully reviewed, discussed and edited the paper. We further thank A. Meyer, D. Dimitrov, M. Magdwick, and K. Agren for helpful discussions regarding algorithmic implementation.

H.M.B. is supported by a MIT-Novo Nordisk Artificial Intelligence Postdoctoral Fellowship and a Cancer Research Institute Immuno-Informatics Postdoctoral Fellowship (CRI12812). NM is supported by the 2024-2025 Takeda Fellowship. D.A.L. is supported by NIH Contract No. 75N93019C00071 and NIH Grant U19-AI135995.

