## Supplementary Information for "scLEMBAS: Context-Aware Modeling of Signaling Pathway Activity at Single-Cell Resolution"

### Supplementary Results and Information

#### Subsetting Perturbations in the McCauley Dataset

Upon quantifying signal strength of each covariate (see Supplementary Methods for details), we observed that the perturbation had the weakest signal strength relative to cell type, cell cycle phase, and batch donor (Fig. S3). Briefly, we fit linear (logistic regression) and non-linear (random forest) models on the TF-activity-derived PCA space using 5-fold cross-validation, assessing model performance using chance-adjusted accuracy, and using model performance as a proxy for covariate signal strength. We reasoned that these results indicate that weak perturbations with sparse transcriptomic effects “wash out” the signal we can reasonably expect scLEMBAS to capture, given that reconstruction error won’t be penalized during training for not accounting for perturbation effects. We note that the limited effect of some perturbations on global, single-cell transcriptomic measurements has been previously observed[1,2], and this is known to reduce single-cell foundation model predictive performance[3].

Consequently,  we subsetted to a set of perturbations labelled as “strong” (i.e., having significant substantial separation from unperturbed cells; see Supplementary Methods for details), which we expected to provide a stronger relative ligand perturbation signal. The five perturbations are TGFB1, IFNG, IFNA2, BMP4, IL13; notably, 4 of these 5 perturbations are listed by McCauley et al. as having the top 5 strongest transcriptional effects by differential expression (all except BMP4). Re-running our latent space embedding and quantification pipeline on this subset of strongly perturbed cells and unperturbed cells, we observed visually (Fig. S3a) and quantitatively (Fig. S3b) that perturbation now had a much stronger relative signal, higher than that of cell cycle phase and batch donor, and comparable to that of cell type (Fig. S3a-b).

To ensure that subsetting to strong perturbations was justified, we also included a couple controls. First, we made sure that the above discrepancy in perturbation signal strength were relevant to training dynamics and attributable to signal washout by weak perturbations on the reconstruction loss, rather than simply having test splits with easier prediction tasks (i.e., only predicting strong perturbations). To do so, we trained models on either all perturbations (“full model”) or strong perturbations only (“subset model”), and had both models predict unseen test data consisting only of strong perturbations. Here, if the full model performs more poorly than the subset model, this indicates that the weak perturbations obscure the ligand perturbation signal such that it is not effectively captured during training. This supports the decision to train *scLEMBAS* on the subset of strong perturbations. In contrast, if the full model performs comparably to that of the subset model, then the perturbation signal can be captured during training, and the above discrepancy in ligand signal strength is simply due to weak perturbations in the test split that are not expected to be captured by predictions regardless. We observed that the full model performed significantly worse than the subset model (Fig. S4a). We also created random subsets of our perturbations that were not categorized as strong, and observed that those models performed significantly worse than the model trained on the strong perturbations (Fig. S4b). This indicates that signal strength discrepancies aren’t simply due to having fewer labels to learn or predict.

#### Model Training Dynamics

Key training dynamic values were tracked across epochs. Firstly, the purpose of the recurrent neural network is to capture feedback loops; consequently, the number of iterations prior to convergence should be substantially larger than the paths between input perturbation and target TFs.  While we didn't implement a regularization on the spectral radius of the recurrent adjacency weight matrix, its value indicates the stability and effective depth of an RNN's dynamics. When the spectral radius is small (<<1), the RNN is strongly contractive, causing hidden states to collapse rapidly to a fixed point and converge in only a few iterations, but with relatively trivial dynamics. As training progresses, optimization can increase the spectral radius, weakening contraction and allowing information to propagate across more steps. When the spectral radius approaches 1, convergence slows substantially and the RNN effectively "uses" its iterative depth, whereas values that surpass 1 lead to instability and divergent or oscillatory behavior. The observed increase in RNN steps over training, alongside spectral radii remaining below the target (0.9), reflects this tradeoff between stability and expressivity within a contractive regime (Fig. S5).

Next, we tracked the various regularization terms and reconstruction loss across epochs, ensuring that reconstruction was the dominant task (Fig. S6). Regularization of the edge sign yielded zero incorrect edge signs when known across all folds (data not shown). Evaluation curves for training followed similar trends (Fig. S6). We also tracked the adversarial training dynamics, demonstrating that the discriminators learn during warmup epochs, and are subsequently fooled through progressive adversarial penalization of the generator, resulting in removal or perturbation and context (cell type) information from the global bias (Fig. S7).

#### Pseudobulk Baseline Comparisons

Given recent work suggesting that single-cell perturbation models fail to outperform—or even underperform—deliberately simple baselines, we benchmarked scLEMBAS against a set of such baselines. Specifically, we emulated the Random Forest (RF) Regressor with GO described in Csendes *et al.*[4], the linear model described in Ahlmann-Eltze *et al.*[5], and the training mean used in both studies. Model performance was evaluated using the same metrics reported in these works: RMSE and Pearson (reported as Pearson delta[6] for the McCauley dataset Pearson correlation correlation for the Kang dataset). Because these baseline models inherently operate on pseudo-bulked perturbation-level data, we applied the same pseudo-bulking to scLEMBAS' predictions. In addition, we evaluated pseudo-bulking by condition (cell type–perturbation combinations), which corresponds to the explicit prediction task scLEMBAS is designed to solve. In this second evaluation, baselines produce identical predictions for a given perturbation across all cell types, reflecting the inherent limitation of the deliberately simple predictive task. This comparison remains aligned with the original intent of these baselines: to establish the performance of deliberately simple predictors; here, we are testing the simple assumption that perturbation relationships are approximately transferable across cell types.

For the McCauley dataset, in the full feature space, scLEMBAS underperformed both the random forest and linear baseline model by both RMSE and Pearson delta when pseudo-bulking by perturbation (Fig. S9b). However, for RMSE, this performance gap disappeared when pseudo-bulking by condition rather than perturbation. This observation is consistent with the design goal of scLEMBAS, which explicitly models cell type–specific perturbation responses rather than perturbation effects alone.

Furthermore, qualitative inspection of condition-specific predictions in the PLS-reduced space suggested that scLEMBAS outperformed the linear and training-mean baselines and achieved performance comparable to the RF baseline (Fig. S10b). Motivated by this observation, we quantitatively re-evaluated model performance in the reduced space (Fig. S11b). In this setting, scLEMBAS significantly outperformed both the linear and training-mean baselines by RMSE when pseudo-bulking by condition and performed comparably to the RF baseline (Cohen's $d$ = 0.1). Furthermore, the worsened performance by Pearson delta disappeared at both the perturbation- and condition- resolution.

Across all twelve comparisons (pseudo-bulking level x model type x performance metric), four Cohen's $d$ effect sizes changed sign when moving from the full feature space to the PLS-reduced space. In three of these cases, scLEMBAS shifted from underperforming to outperforming the corresponding baseline. Among the remaining eight comparisons, five of six cases in which scLEMBAS underperformed showed reduced effect sizes, whereas both cases in which scLEMBAS outperformed a baseline showed increased effect sizes. Together, these results indicate that dimensionality reduction reveals improved relative performance of scLEMBAS compared to pseudo-bulk baselines.

For the Kang dataset, results are qualitatively similar, though more apparently outperforming baselines(Fig. S9a, Fig. S10a, Fig. S11a): scLEMBAS is not outperformed by baselines when considering perturbations, and it significantly outperforms the baselines when considering conditions. However, in this instance, it significantly outperforms baselines by both RMSE and Pearson correlation and even in the full feature space. While significance is consistent when considering PC space, effect sizes are improved as with the McCauley dataset: Across all twelve comparisons, four Cohen's $d$ effect sizes changed sign when moving from the full feature space to the PC space. In all of these cases, scLEMBAS shifted from underperforming to outperforming the corresponding baseline. Among the remaining eight comparisons, both cases in which scLEMBAS underperformed showed reduced effect sizes, whereas all six cases in which scLEMBAS outperformed a baseline showed increased effect sizes.

Overall, these results highlight two key takeaways: each of accounting for cell type and using a reduced space improves the relative performance of scLEMBAS. When considered jointly, scLEMBAS tends to outperform baselines by RMSE for the McCauley dataset; moreover, scLEMBAS outperforms baselines in the full feature space by both metrics for the Kang dataset when accounting for cell type.

#### Global Assessment of Counterfactual Perturbation Predictions

In addition to univariate differential feature analysis to determine whether predicted perturbations are consistently separated from their corresponding controls, we also used the Mahanalobis distance as a global metric that accounts for all features simultaneously. In contrast to differential feature analysis, which assessed separation between predicted counterfactuals and actual data control unperturbed cells, here we specifically quantified the Mahanalobis distance between predicted counterfactuals and predicted controls. In other words, the conditions in test splits underwent the standard counterfactual forward pass, and are compared to the corresponding condition (same cell type, control perturbation) in the train split that underwent the forward pass without a counterfactual as applied during training (both are extensively described in the Methods). This analysis isolates a failure mode in which predictions may appear globally accurate yet fail to distinguish perturbation effects from control, with the model failing to learn true perturbation separation. Such a failure mode could reflect incomplete adversarial removal, leakage of perturbation-relevant information through other architectural components, memorization rather than generalization, mode collapse[7], or averaging of heterogeneous perturbation responses under squared-error reconstruction losses[8].

For the McCauley dataset, for each test condition, we learned a latent space defined by the perturbation and its corresponding control of the same cell type using partial least squares separating perturbation status, ensuring that the latent space emphasizes counterfactual-specific variation.Upon quantification, we observed that scLEMBAS captures meaningful perturbation–control separation compared to the baseline model that received no adversarial penalization, as reflected by an OLS coefficient closer to 1 (Fig. S13a). This indicates higher agreement with the identity line, representing perfect predictions, and is supported by a significantly lower residual distance to the identity line with a large effect size (Fig. S13b). Because this baseline differs only in the absence of adversarial penalization, relatively improved separation between predicted counterfactuals and predicted controls reflects scLEMBAS' ability to learn counterfactual structure beyond what is required to optimize predictive accuracy as defined by the loss function.

We note, however, that the predicted counterfactual separation is reduced relative to the actual data, reflected by the OLS coefficient being less than 1. This is likely associated with the aforementioned failure modes. However, it learns sufficiently well enough to capture information regarding univariate differential features (Fig. 2c, Fig. S14).

#### Subtype Perturbation Response Geometries

Amongst cell subtypes, we categorized perturbation-induced expression shifts[9] into three response categories : (1) divergent, in which subtype centroids are further apart upon perturbation, convergent, in which subtype centroids are closer together upon perturbation, or parallel, in which subtype centroids shift in a parallel manner upon perturbation (Fig. S16a). The category of expression shift must be considered due to the nature of the counterfactual, in which the origin perturbation is provided to the model via the gene expression vector. Thus, we aim to ensure that the subtype heterogeneity of the target perturbation is preserved regardless of the expression shift category.  We note that, under the typical counterfactual—transitioning from a control cell to the same cell type under perturbation—expression shifts were generally convergent or parallel. Consequently, we also evaluated counterfactuals exhibiting divergent shifts, in which subtype centroids in the origin perturbation (i.e., the model gene expression input) were closer together than those in the target perturbation (see Methods for details).

Upon ensuring all these geometries were represented, we quantified the ability of scLEMBAS to capture subtype specific responses using the Mahanalobis distance (for subtype expression shift magnitude) and cosine distance (for subtype expression shift geometry) (Fig. 2d-e). Since each cell type and perturbation pair assessed (scatter points) represents one of three expression shift types, we are capturing a range of expression shift types; thus, the globally positive relationship between predicted and observed separation across both metrics indicates that scLEMBAS captures subtype-specific perturbation heterogeneity irrespective of expression shift type. In other words, subtype heterogeneity is preserved with a consistent geometry across varying magnitudes of target perturbation subtype separation as well as varying relative magnitudes of origin perturbation subtype separation.

#### Edge Weight Regularization Hyperparameters Must be Altered to Observe Self-Pruning

Ensemble training was consistent with other models trained on the McCauley dataset, with one notable exception: the L2 regularization of the signaling network weights across a range of values up to 7 orders of magnitude larger than that typically used did not result in self-pruning. Instead, we implemented L1 regularization to achieve self-pruning. The regularization coefficient was set to 1e-3, with values one order of magnitude lower than this also failing to produce this effect. We confirmed that, at this level of regularization and with the inclusion of spurious edges, the ensemble models' predictive performance is not significantly different than that of the standard models used in our 5-fold CV assessments (Fig. S18a). We also observed qualitatively similar training dynamics across epochs (compare Fig. S7b for the standard models using L2 regularization to Fig. S18b for the ensemble models with spurious edges using L1 regularization).

#### Large Magnitude Spurious Edges Have an Outsized Effect on AAD

Carrying the discussed GAMM analysis further, when removing edges, we removed either spurious edges or an equal number of real edges; we also fit our GAMM while controlling for the edge type. Consequently, we could specifically determine the effect of edge type on scLEMBAS' predictive capacity. When including a weight-type interaction in our GAMM (likelihood ratio test $\chi^2$ $p < 2.2e\text{-}16$)–with the reduced model defined as one that does not include the interaction–the AAD of real edges was greater than that of false edges across a majority of weights (Fig. S17b). However, notably, at the ~$75^{th}$ percentile of weight magnitudes, spurious interactions begin to have a greater AAD than real interactions. Together with our self-pruning results in this percentile range (Fig. 3e-f), this tells us that while scLEMBAS tends to downweight spurious interactions in the high-weight magnitude range, in instances where large spurious interaction weight magnitudes are retained, they have a substantial impact on model prediction. Intuitively, if the model seems to be identifying spurious interactions overall, but weighs some heavily, it makes sense that these edges matter to the model. These may be due to "shortcuts" yielding simpler paths between perturbation input and TF activity output or genuine missing interactions in the PKN, meriting further future investigation. These results are not consistent with that of the bulk version of LEMBAS[10], in which AAD values of real edges were consistently greater than or equal to that of spurious edges across the range of learned magnitude weights.

### Supplementary Methods

#### Latent Space Covariate Signal Quantification

Upon embedding data into a latent space, we quantified how much biological and technical covariate information (from the observation metadata) is captured by the latent space. In particular, for a given covariate, we regressed each covariate of interest onto all retained latent variables jointly using stratified 5-fold cross-validated classifier prediction. To capture both the linear and nonlinear structure, we used a logistic regression and random forest classifier, respectively. Logistic regression was fit using *scikit-learn* (saga solver, L2 penalty; max_iter = 1000). Features were standardized to zero mean and unit variance prior to model fitting to aid convergence. Random forest was fit using *scikit-learn*'s default features. These were also the same classifiers used for assessing the extent of adversarial removal in the global bias. The information captured by the latent space was quantified by the classifier chance adjusted accuracy, enabling comparison across covariates. The chance-adjusted accuracy metric normalizes classification accuracy relative to the expected accuracy under random guessing, assuming equal class probability. This score equals 0 at chance performance and 1 at perfect classification. This assessment is particularly informative with PCA embeddings, which capture maximal directions of variance in a fully unsupervised, covariate-agnostic manner.

#### Labelling McCauley Perturbations as “Strong”

Pairwise E-distances between perturbation conditions were computed in PCA space using the “edist” function from the Python package *scperturb*[11] with default values. For each perturbation, a global separation score was defined as the median pairwise distance to all other perturbations (excluding self-comparisons). Perturbations were categorized as "strong" if they had a significant difference from the control condition. To assess significance, we used a bootstrap procedure to estimate the 95% confidence interval of the difference in global separation scores between each perturbation and the control. Specifically, for each of 5,000 bootstrap iterations, global separation scores for the perturbation and control were estimated by resampling each median distance distribution with replacement and taking the difference. Perturbations with a two-sided confidence interval lower bound greater than 0 were categorized as significantly different from control.

#### Controls for McCauley Perturbation Subsetting

We ran two control comparisons to further probe our choice in subsetting to strong perturbations. Since these involved subsets of the data and direct comparisons, we fit models on the full feature space rather than principal component space. For a given comparison, we fit two model types: linear and non-linear as described in the “Latent Space Covariate Signal Quantification” section. To accommodate the higher-dimensional feature space relative to PC space, logistic regression was fit using the lbfgs solver with increased regularization (C = 0.5) and an increased maximum iteration limit (max_iter = 3000). Feature standardization was not applied since consensus transcription factor activity scores are already z-scored.

In the first control, we compared models fit on all perturbations (“full model”) to those fit on strong perturbations only (“subset model”). To do so, we created stratified 10-fold splits on all perturbations. Next, we subsetted the splits to strong perturbations only (“subset train” and “subset test”). Finally, we downsampled the train split to match the sample size of the strong perturbation subset (“full train”). We ensured that all subsetted and downsampled splits preserved corresponding class frequencies within a tolerance of 1 percentage point. The full and subset models were fitted on the full and subset train splits, respectively, and both were assessed on the subset test split using the balanced accuracy score. This ensured that both models had equal sample sizes when fit, and frequencies that

In the second control, we compared models fit on weak perturbations to models fit on the strong perturbations. For weak perturbations, we created 10 random subsets of perturbation labels that were not included in the strong perturbations set. Each random subset satisfied all following conditions:

1. The number of perturbation labels equalled that of the strong perturbation set (6 total labels)
2. The number of cells that fall into the weak set perturbation labels is approximately equal to that of the strong perturbation labels (within a tolerance range of 10% of the number of cells in the strong perturbation labels)

3. The rank-ordered distribution of perturbation label frequencies approximately matched that of the strong perturbation set, such that the frequency of the most abundant weak label matched that of the most abundant strong label, the second most abundant weak label matched the second most abundant strong label, and so on, within a tolerance of 2.5 percentage points.

Next, for both the strong perturbation set and each weak perturbation set, we ran stratified 5-fold cross-validation and assessed model performance using the balanced accuracy score.

#### Pseudobulk Baseline Comparisons

We used three pseudobulked baseline models: (1) Random Forest (RF) Regressor with GO described in Csendes et al.[4], (2) the linear model described in Ahlmann-Eltze et al.[5], and (3) the training mean used in both studies. Pseudobulking is conducted by taking the mean across cells for a given label.

For the RF baseline, the GO perturbation embedding was generated following Csendes *et al.* Specifically, we downloaded their binary gene–Gene Ontology indication matrix (https://github.com/turbine-ai/PerturbSeqPredBenchmark/blob/main/data/go/go_raw_matched.csv) and applied principal component analysis (PCA) to 256 components. To include control perturbations, we added an all-zero control vector to the indication matrix prior to PCA. Because the RF input features depend only on the perturbation embedding and are therefore identical for all cells receiving the same perturbation, we did not apply explicit pseudo-bulking prior to model fitting; the model effectively pseudo-bulks implicitly, producing the same predicted output for identical perturbation features.

For the linear baseline, we calculated the feature embedding `G` by applying PCA to the perturbation-level pseudo-bulked training data. PCA was performed to two components for the McCauley dataset and to one component for the Kang dataset (which contains only two perturbations). Because `G` is computed from transcription factor (TF) activity rather than gene expression, it cannot be used to recover the perturbation embedding `P` as defined in Ahlmann-Eltze *et al.* Instead, we used the same GO-based perturbation embedding employed for the RF baseline.

We applied the linear baseline to match scLEMBAS' prediction task. While Ahlmann-Eltze *et al.* used their linear model to predict responses to unseen perturbations, scLEMBAS predicts unseen cell type–perturbation combinations while all individual perturbations and cell types have been observed during training (analogous to the compositional perturbation autoencoder (CPA)[12]). Ahlmann-Eltze *et al.* used CPA for predicting unseen perturbations and acknowledge this is an unfair task not directly comparable to the baseline. Here, we instead reverse this logic: at test time, the linear model reuses the perturbation embeddings `P` observed during training. Consequently, the model applies the learned linear transformation matrix `W`—which encodes feature-perturbation relationships from training—directly to test data without any adaptation to unseen perturbations. This baseline therefore tests whether perturbation effects learned during training transfer directly to new cellular contexts, analogous to the training-mean baseline but with perturbation-specific structure retained.

To assess model performance, we evaluate predictions using root mean squared error (RMSE) and Pearson-based metrics, following Ahlmann-Eltze *et al.* For the McCauley dataset, we apply the Pearson delta as described in Cui *et al.*[6] and employed by both Ahlmann-Eltze *et al.*[5] and Csendes *et al.*[4]. The Pearson delta requires a control perturbation, which assumes that all test predictions are non-control. Because the Kang dataset train–test split includes both control and perturbed conditions in the test set, we instead use the Pearson correlation directly for simplicity.

All metrics were computed sample-wise (i.e., on pseudo-bulked labels), and the reported values correspond to the mean across samples. Metrics were evaluated both in the full feature space and in a reduced latent space. For the McCauley dataset, the reduced space for each fold was obtained via partial least squares (PLS), following the described latent space embedding pipeline, applied to the test data using test condition labels (cell type–perturbation combinations) as the response. For fold 4, visual inspection of the 5-fold cross-validation accuracy as a function of the number of PLS components revealed a non-monotonic curve with multiple local elbows. While the automated elbow criterion selected two components, accuracy continued to improve gradually with additional components. We therefore manually selected 10 components for this fold to capture the broader performance plateau rather than a local elbow. We ensured all model fits are significant (100 permutations Q2Y null p-value ≤ 0.05). For the Kang dataset, the previously described PC space fit on all the actual data was used. Model predictions were projected into the latent space.

#### Categorical Bias: Subnetwork Identification

To assess the relevance of the categorical bias, we focused on club and basal cells in the McCauley dataset under TGFB1 perturbation. This represented a pair of cell types with sufficient cell numbers (>75) in both control and TGFB1-perturbed conditions for each cell type. Furthermore, the cell types demonstrated distinct expression shifts and perturbed states in response to TGFB1 (Fig. 4d). Our ultimate goal was to identify the bias nodes most relevant to distinguishing the cell-type-specific perturbation responses.

As a preliminary filter, we identified a subnetwork of protein-protein interactions that defined the club and basal cell TGFB1 response. First, we retained only the top 5 TFs, rank ordered by the PLS fit on this subset of data, using a multivariate response defined by both cell type (club or basal) and perturbation (control or TGFB1). Next, to identify signaling interactions supporting the TGFB1 response in both cell types, we implemented a single-cell, ensemble adaptation of a previously developed gradient-based subnetwork extraction framework[13]. The procedure was applied independently to 10 ensemble models from each fold of the five-fold cross-validation, corresponding to 50 trained models in total. For the edge-importance calculation, equal numbers of unperturbed basal and club cells were sampled, with a maximum of 200 cells per cell type. When the two populations differed in size, the larger population was subsampled without replacement using a fixed random seed. Each cell was retained as an individual context, preserving its expression input and cell-type covariate. We then defined a uniform TGFB1 stimulation path using 21 evenly spaced nonzero input perturbation levels, $s_m = m/21$ for $m = \{1, 2,..., 21\}$, with a zero-input forward pass serving as the corresponding control.

For each trained model, we accumulated the baseline-adjusted sensitivity of the predicted TF response to each signaling edge across this stimulation path. Specifically, we summed the TGFB1-induced change across all modeled TF outputs, stimulation levels, cells, and the two cell types and differentiated this aggregate response with respect to each learned signaling weight. The importance of edge $e$, parameterized by $w_e$, was defined as:

$$I_e = \left| \frac{\partial}{\partial w_e} \left[ \sum_c \sum_{i=1}^{n} \sum_{m=1}^{21} \sum_{k=1}^{K} (\hat{y}_{cimk} - \hat{y}_{ci0k}) \right] \right|,$$

Where c is one of the two {basal cell, club cell}, $\hat{y}_{cimk}$ denotes the predicted activity of modeled output TF $k$ for cell i of cell type $c$ at TGFB1 input level $s_m$, and $\hat{y}_{ci0k}$ denotes the corresponding prediction at zero TGFB1 input. This forms a Riemann approximation of integrated parameter sensitivity along the TGFB1 stimulation path. We therefore refer to it as a customized integrated-gradient score. The path was defined over TGFB1 input, whereas gradients were calculated with respect to signaling weights. Scores for positions absent from the prior knowledge network and for incoming interactions to the TGFB1 input node were set to zero. Edge importance was calculated using all modeled TF outputs, whereas the subsequent fidelity criterion was evaluated using the five prespecified TFs of interest.

For each model, we evaluated 200 logarithmically spaced importance thresholds spanning the model-specific range of positive scores, together with a threshold of zero. At threshold $\tau$, signaling weights associated with edges of importance below $\tau$ were set to zero, and control-to-TGFB1 counterfactual predictions were recomputed at unit TGFB1 input using all available unperturbed basal and club cells. For each cell type, fidelity to the complete model was quantified using the mean absolute error (MAE) over cells and the TFs of interest, normalized to the MAE obtained when all signaling network weights were set to zero:

$$E_c(\tau) = \frac{\mathrm{MAE}\left(\hat{\mathbf{Y}}_c^{(\tau)}, \hat{\mathbf{Y}}_c^{\mathrm{full}}\right)}{\mathrm{MAE}\left(\hat{\mathbf{Y}}_c^{\mathrm{zero}}, \hat{\mathbf{Y}}_c^{\mathrm{full}}\right) + 10^{-12}}, \qquad E_{\mathrm{TOI}}(\tau) = \frac{E_{\mathrm{Basal}}(\tau) + E_{\mathrm{Club}}(\tau)}{2}.$$

We selected the largest threshold satisfying $E_{TOI} \leq 0.25$, corresponding to the most stringent pruning level that retained the specified prediction fidelity (Fig. S20). If no threshold satisfied this criterion, the threshold minimizing $E_{TOI}$ was retained. Mean TF-wise Pearson correlations, the number of reachable TFs, and the number of directed TGFB1-to-TF paths were recorded (together with a few other metrics) as diagnostics but did not constrain threshold selection (Fig. S20).

The directed graph for each model retained edges whose importance exceeded the selected threshold. We removed incoming edges to TGFB1, and TGFB1 outgoing edges that did not terminate at a candidate receptor. For TGFB1, candidate receptor representations comprised the TGFBR1_TGFBR2 complex and the individual TGFBR1 and TGFBR2 nodes when present in the model. Each surviving receptor r was scored from all simple paths of at most eight edges connecting it to the TFs of interest. For path p, path importance was defined as the geometric mean of its edge-importance scores: $Q(p) = \left( \prod_{e \in p} I_e \right)^{1/|p|}$. Receptor scores were calculated by first averaging Q(p) across paths to each reachable TF and then averaging across reachable TFs. The highest-scoring receptor determined whether the complex or individual-receptor representation was retained. If no receptor had a positive path score, all surviving receptor candidates were retained. Finally, each per-model graph was restricted to the intersection of nodes reachable from TGFB1 and nodes capable of reaching at least one TF of interest. Therefore, every retained node lies on at least one directed TGFB1-to-TF path.

Per-model subnetworks were aggregated by directed source–target edge identity. For each edge, we recorded its frequency of retention and calculated its mean learned weight and mean importance across the models in which it was retained. Consensus frequencies across all folds were normalized by the number of successfully processed models. The consensus subnetwork retained edges present in at least 50% of these models. Edge sign was assigned from the sign of the mean learned weight, and the consensus graph was again restricted to nodes lying on a directed path from TGFB1 to at least one TF of interest. All subsequent node-level analyses were restricted to this consensus subnetwork .

All subsequent analyses with the categorical bias used only the subnetwork identified here.

#### Categorical Bias: Node Importance

Nodes in the subnetwork were rank-ordered using the categorical bias term, emulating a previously described approach[13] with some modifications. Node importance scores were calculated for every signaling node in each trained model using the same balanced basal and club control cell sets and the same 21-level TGFB1 stimulation grid. Conceptually, for each cell and stimulation level, we considered both the magnitude of the bias term and how it's incorporated with the remaining model components. Specifically, with regards to its incorporation with remaining model components, we calculated the absolute change in the steady-state activity of every signaling node between the TGFB1-stimulated and zero-input forward passes. These activities were obtained directly from the signaling network before application of the TF output layer and after iterating through the RNN.

For signaling node j, the implemented importance score was:

$$N_j = \frac{1}{2} \sum_{c \in \{basal,\ club\}} \left| b^{cat}_{c,j} \right| \left| Y_{c,i,m,j} - Y_{c,i,0,j} \right|$$

where i indexes cells, m indexes TGFB1 input levels, *Y* represents the steady-state activity of signaling nodes that is output by the RNN, and $b^{cat}_c$ is the categorical bias of cell type *c*. Therefore, the implemented score combines the magnitude of the categorical bias with the perturbation-induced change in node activity. Scores represent averaging across basal and club cells. Nodes were rank-ordered according to this final value . We repeated this for each ensemble model and calculated the median rank, giving us our final node importance score. These ranks were subsequently intersected with the consensus-subnetwork nodes for downstream analyses.

Node importance was compared to the absolute value of the $\log_2$-fold-change (LFC) effect size output from differential expression (DE) analysis between club and basal cells. DE was conducted using an orthogonal dataset, the Human Lung Cell Atlas (HLCA v1.0)[14]. Data was filtered to exclude donors below 24 years of age and above 54 years of age, as well as club cells annotated as “nasal”. Raw count data were aggregated by summing counts across cells within each unique sample-by-cell-type combination using decoupler's pseudobulk function. To ensure adequate representation, pseudobulk profiles were retained only if they were derived from at least 10 cells and contained a minimum of 1,000 total counts. The resulting count matrix was analyzed with PyDESeq2. A DESeq2 model was fit with cell type as the design factor, using Cook's distance–based refitting of outliers. Differential expression was tested for the contrast comparing club cells against respiratory basal cells, with basal cells serving as the reference level. Single-threaded inference was used throughout. Genes were considered significantly differentially expressed if they met both an adjusted p-value threshold (Benjamini–Hochberg adjusted p-value < 0.05) and an effect-size threshold of |LFC| > 1.5. Significant genes were then ranked by the absolute magnitude of |LFC|.

#### Categorical Bias: Candidate Node Alteration

To identify candidate perturbation targets, we focused on the set of significant genes that were not TFs or ligand perturbations in the McCauley dataset. For each candidate node, three factors were considered: a node importance, the absolute LFC effect size from the differential expression results, and network centrality quantified by PageRank on the full network (not the subnetwork).

Each factor was independently rescaled to a common (0, 1] range based on each node's relative standing within that feature, so the three features could be combined directly. To allow each feature to contribute in the intended direction, inverted versions of the centrality and absolute-fold-change percentiles were also computed (1 minus the percentile). A composite priority score was then formed as a weighted sum of these normalized features, with 60% weight on the node-ranking signal as the primary criterion, 20% on the inverted absolute fold change (to upweight candidates overlooked by differential expression), and 20% on inverted centrality (to downweight candidates with likely pleiotropic effects).

For a given subset of nodes (identified as the top n by the candidate node composite priority score), scLEMBAS model parameters were altered as follows: one cell types categorical bias values were replaced by a scaled (between 1x to 1e4x) of the other. Next, model forward passes were re-run with the new parameters, predicting the altered cell type under TGFB1 perturbation. Forward pass predictions only included ensemble models in which the perturbed cell type was present in the training data without a counterfactual, as the goal here was not to evaluate out-of-distribution prediction, but to use learned parameters to alter cell state.

Finally, two metrics–cosine distance and euclidean distance–were used to assess the extent of “recovery”, i.e., the extent to which the altered cell type’s perturbation state now reflected that of the corresponding cell type. The euclidean distance was calculated simply as the distance of the altered cell type prediction centroid to the centroid of the corresponding cell type derived from the actual data under TGFB1 prediction. The cosine distance was taken between two vectors, each originating at the centroid of unperturbed club and basal cells. One vector terminated at the centroid of predictions for the altered cell type, and the other at the the centroid of the corresponding cell type derived from the actual data under TGFB1 prediction. Cosine and Euclidean distances were each converted to similarity scores, by subtracting the cosine distance from one and by taking the inverse of one plus the Euclidean distance, respectively.

### Supplementary Figures

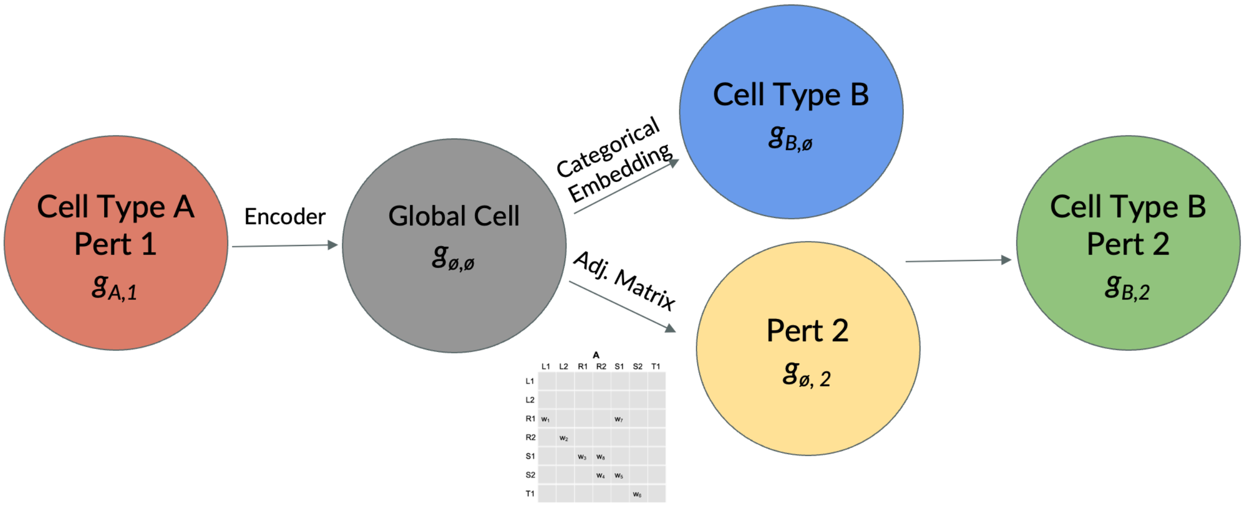


**Figure S1: Flow chart outlining how each of scLEMBAS' architecture components contributes to OOD prediction**. Given an input gene expression vector of a cell representing "Cell Type A" and "Perturbation 1", scLEMBAS is capable of asking the counterfactual question: What would the TF activity of this cell have looked like as "Cell Type B" and/or under "Perturbation 2". This requires both "Cell Type B" and "Perturbation 2" to have been seen in training, but their combination has not been measured. The stochastic encoder removes "Cell Type A" and "Perturbation 1" information from the input gene expression. The categorical bias adds "Cell Type B" information. The adjacency matrix encodes the propagation of information from Perturbation 2 represented by "***x***" while learning the relative interaction strength of the network. Note, most predictive counterfactual tasks in this manuscript consider the same cell type, from the control unperturbed condition to a perturbation.

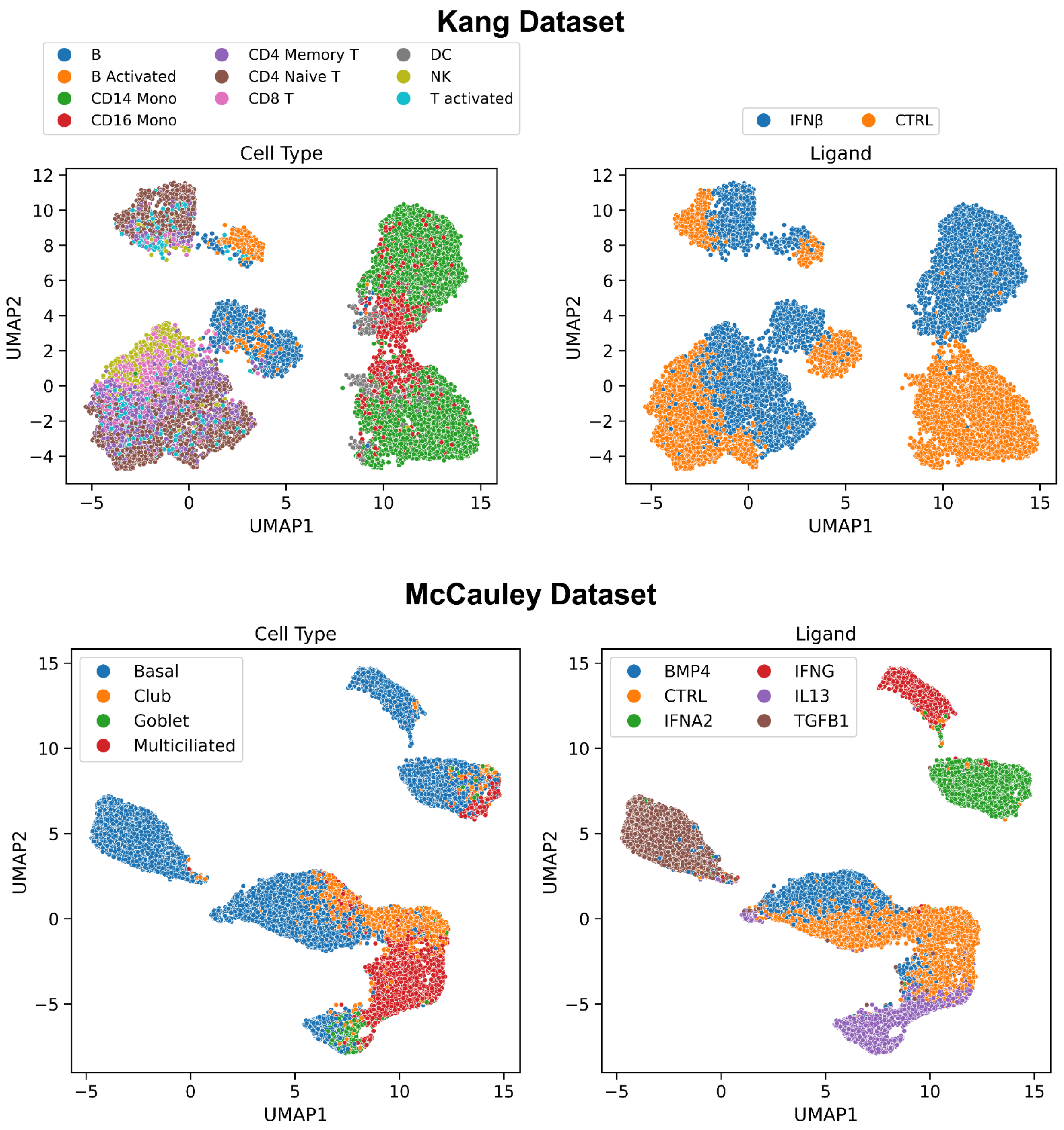


**Figure S2:** UMAPs of TF activity inferred from single-cell RNA-sequencing for the Kang (top panels) and McCauley (bottom panels) datasets. UMAPs were fit on PCA and PLS (maximizing perturbation separation) latent spaces for the Kang and McCauley datasets, respectively. PCA/PLS and downstream UMAPs were fit after all preprocessing was implemented, including signaling network filters.

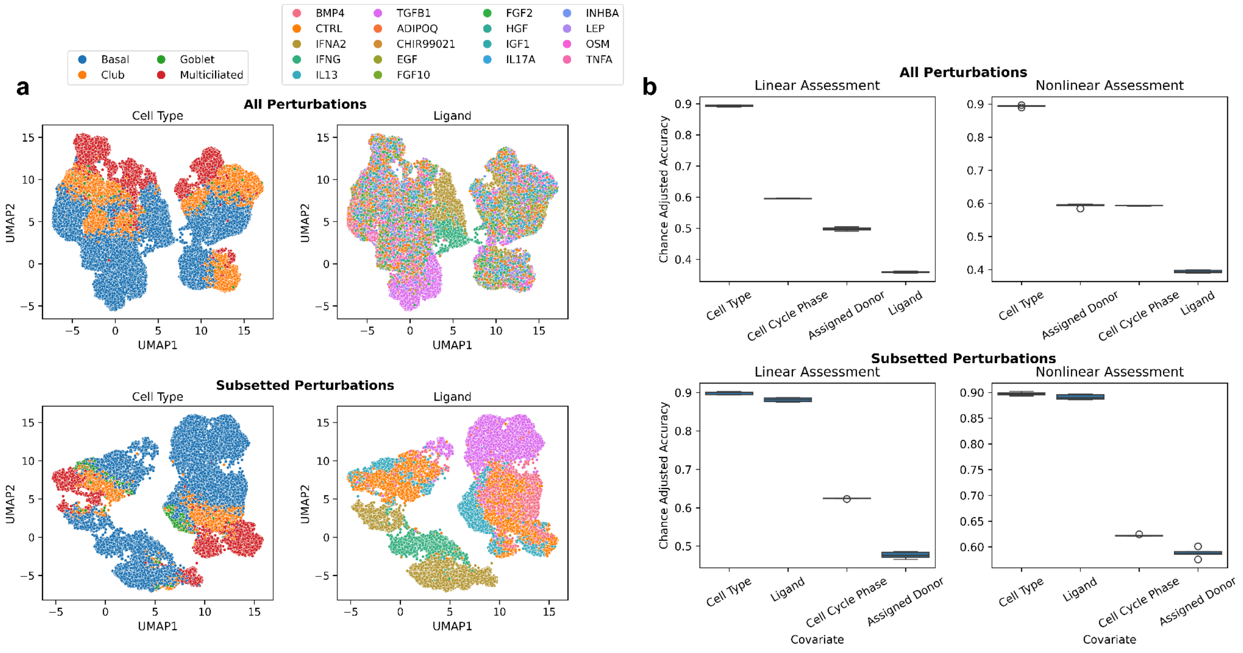


**Figure S3**: **Subsetting in the McCauley dataset increases perturbation-driven signal. (a)** UMAPs of TF activity inferred from single-cell RNA-sequencing for McCauley dataset for all perturbations (top panels) and subsetted to strong perturbations only (bottom panels). UMAPs were fit on PCA latent spaces. **(b)** Boxplots display resultant 5-fold cross-validation test chance-adjusted accuracy scores (y-axis) for each covariate (x-axis) from the signal quantification pipeline. Chance-adjusted accuracy was assessed using linear (logistic regression, left panels) and non-linear (random forest, right panels) models. Quantification was assessed on PCA space fit on each of the full dataset (top panels) and subsetted to strong perturbations only (bottom panels).

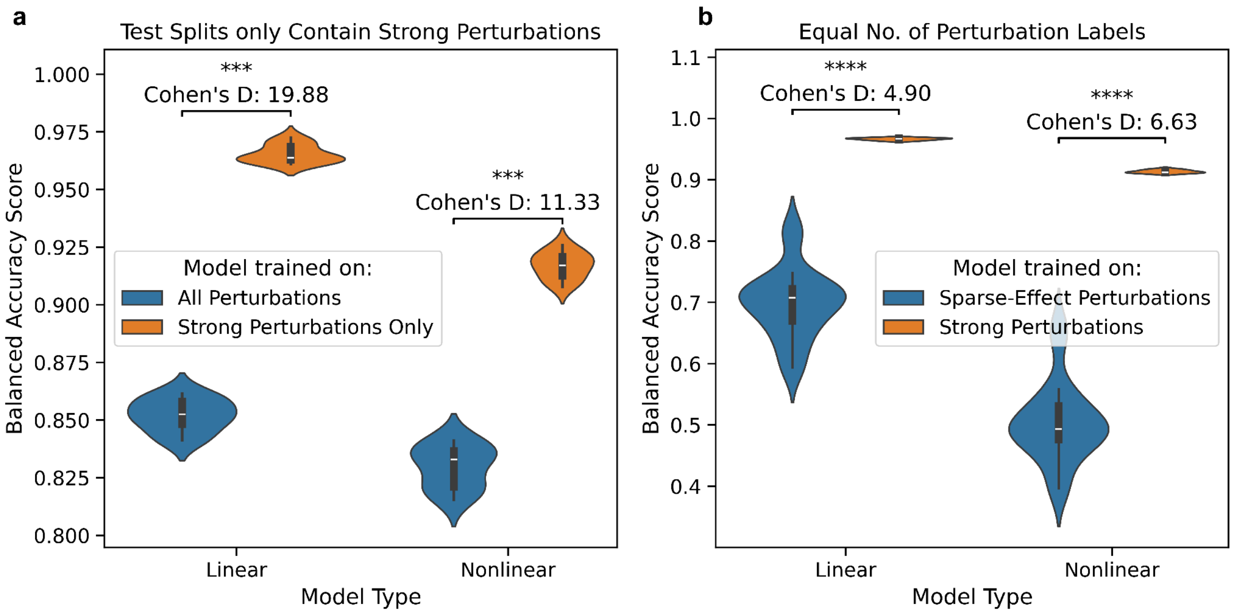


**Figure S4: Control Models Assessing Perturbation Subsetting**. Violin plots display cross-fold validation test balanced accuracy scores (y-axis) for linear (logistic regression) and non-linear (random forest) classifiers (x-axis). Comparisons are quantified by Cohen's D effect size and assigned significance using a Mann-Whitney U significance test (p≤5e-5 ****, p≤5e-4 ***, p≤5e-3 **, p≤5e-2 *). **(a)** 10-fold CV of scLEMBAS models trained on either all perturbations or strong perturbations only, each assessed on test splits consisting of strong perturbations only. Test splits were identical across models, and training splits were matched in sample size. **(b)** 5-fold CV of models trained on strong perturbations only or a subset of sparse-effect perturbations that were not categorized as strong. Sparse-effect perturbation subsets were matched with that of the strong perturbations for number of labels, approximate sample size, and approximate label frequency distribution.

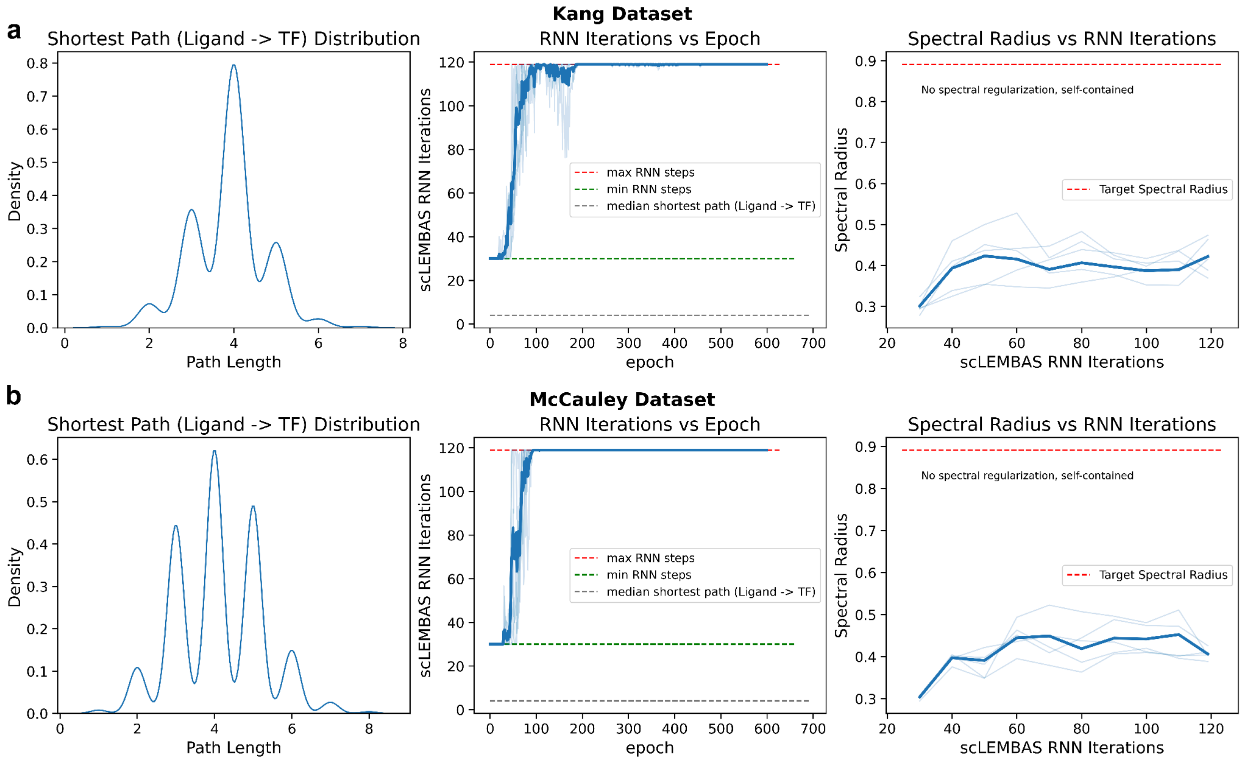


**Figure S5: Recurrent neural network training dynamics.** For each of the **(a)** Kang and **(b)** McCauley datasets, we visualize the distribution of shortest path lengths between input perturbation and target TF (left panel), the number of iterations in the recurrent neural network as a function of epoch (middle panel), and the value of the adjacency matrix spectral radius as a function of the number of RNN iterations (right panel). The bold line displays the average across models, whereas the lighter lines display results for each individual model in 5-fold CV.

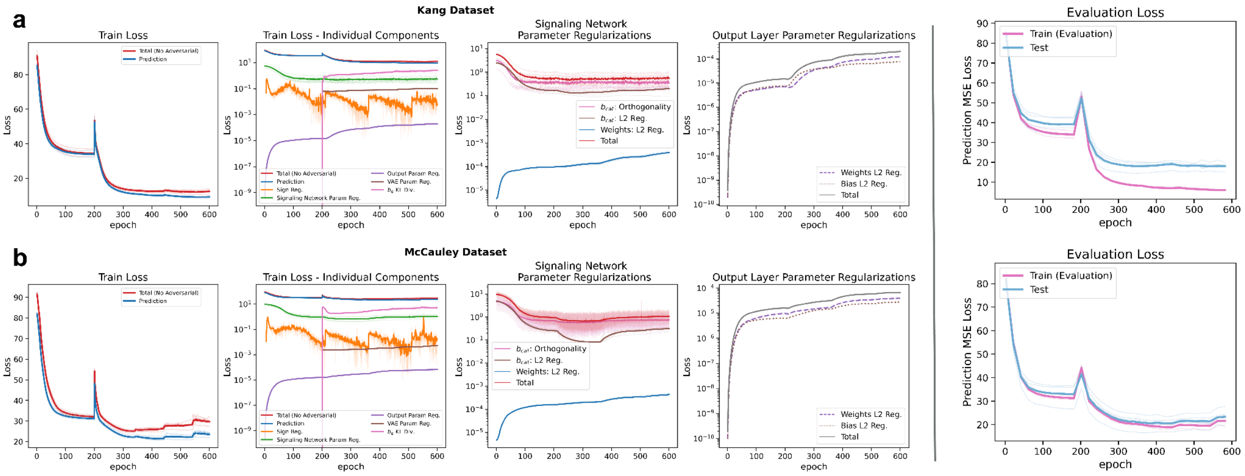


**Figure S6: Loss and regularization training dynamics.** Training loss and regularization values across epochs for scLEMBAS models. The bold line displays the average across models, whereas the lighter lines display results for each individual model. Evaluation loss curves include OOD test predictions (blue line) and train loss evaluated without training-specific effects (e.g., regularizations and noise injection; pink line).

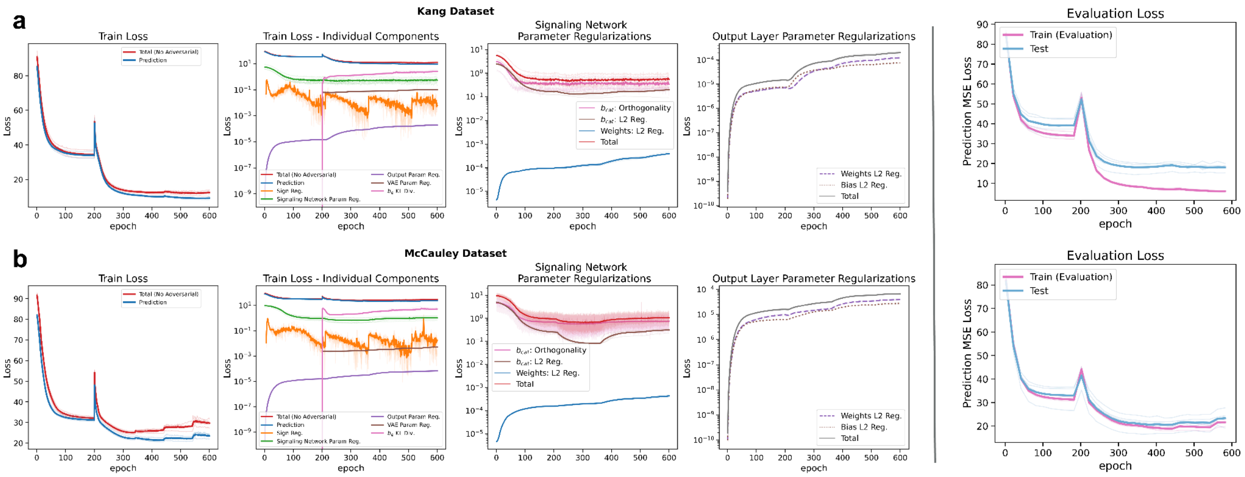


**Figure S7: Adversarial training dynamics**. Training loss decomposed by adversarial penalizations (left panels) and discriminator losses (middle and right panels)  across epochs for models. The bold line displays the average across models, whereas the lighter lines display results for each individual model. Discriminator parameters were regularized with L2 penalties for the Kang dataset, whereas spectral normalization was applied for the McCauley dataset. Horizontal gray dashed lines represent random prediction values for the discriminator.

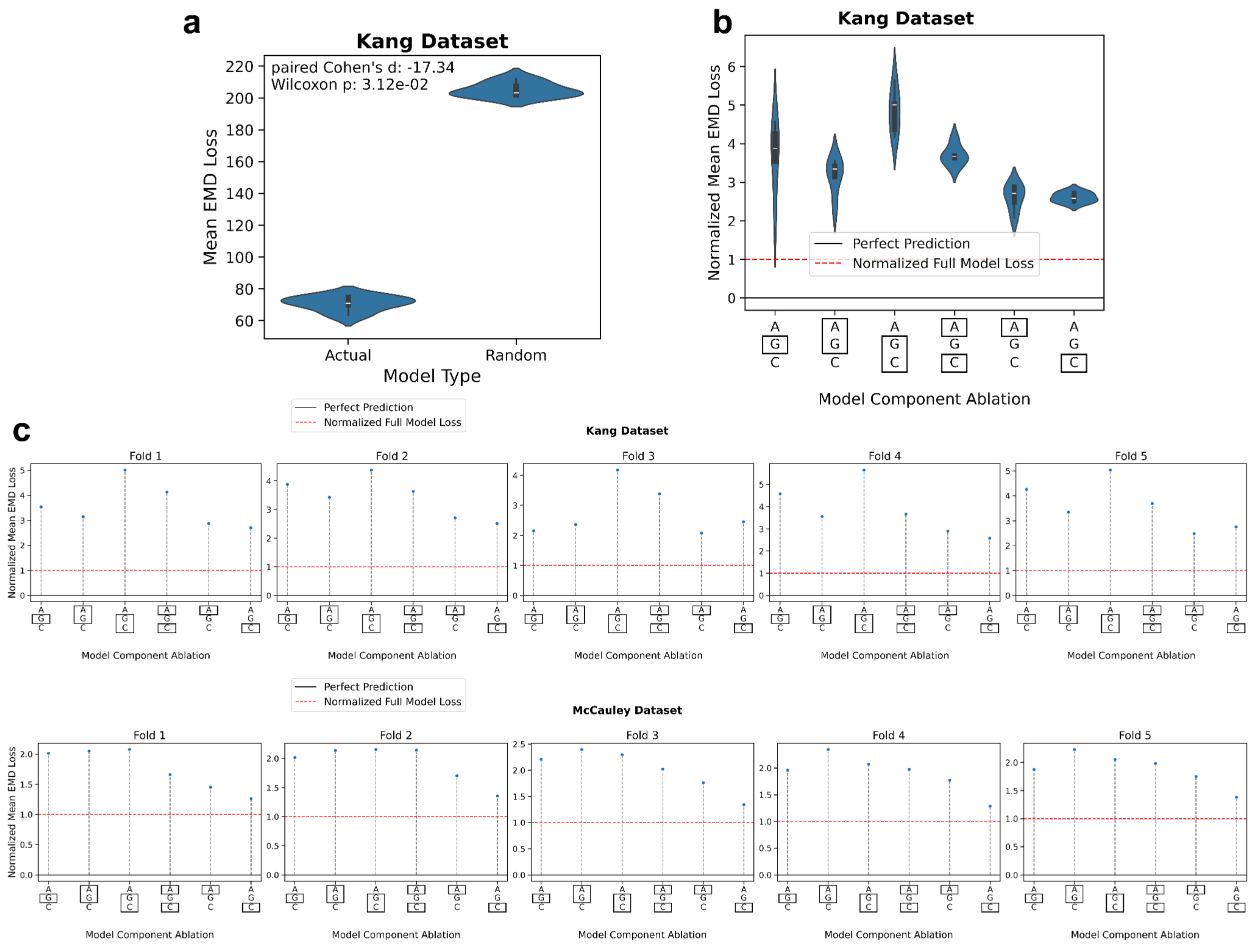


**Figure S8: Change in EMD loss upon model component ablation. (a)** Violin plots compare the distribution of EMD loss between the actual model and the random baseline model. For each fold, loss was calculated separately for each individual test condition, and the mean across conditions was taken. This is the same comparison as the EMD loss in Fig. 1c, but for the Kang dataset. **(b)** Violin plots showing the across-fold distribution of normalized EMD loss for models with different component ablations. Within each fold, EMD loss is computed for each test condition and averaged across conditions. For each ablated model forward pass, losses are normalized to the corresponding full scLEMBAS model (red dashed line), such that values greater than one indicate worsened predictive performance (i.e., a positive contribution of the omitted component to model accuracy). X-axis labels denote the ablated component with a black box, with unboxed components retained in the forward pass. “A” represents the adjacency matrix, “G” the global bias, and “C” the categorical bias. **(c)** Lollipop plots showing the normalized EMD loss for models with different component ablations for individual folds for the Kang (top panel) and McCauley (bottom panel) datasets. This shows the individual fold values creating the violin plot distributions in panel (b) here and Fig. 1e.

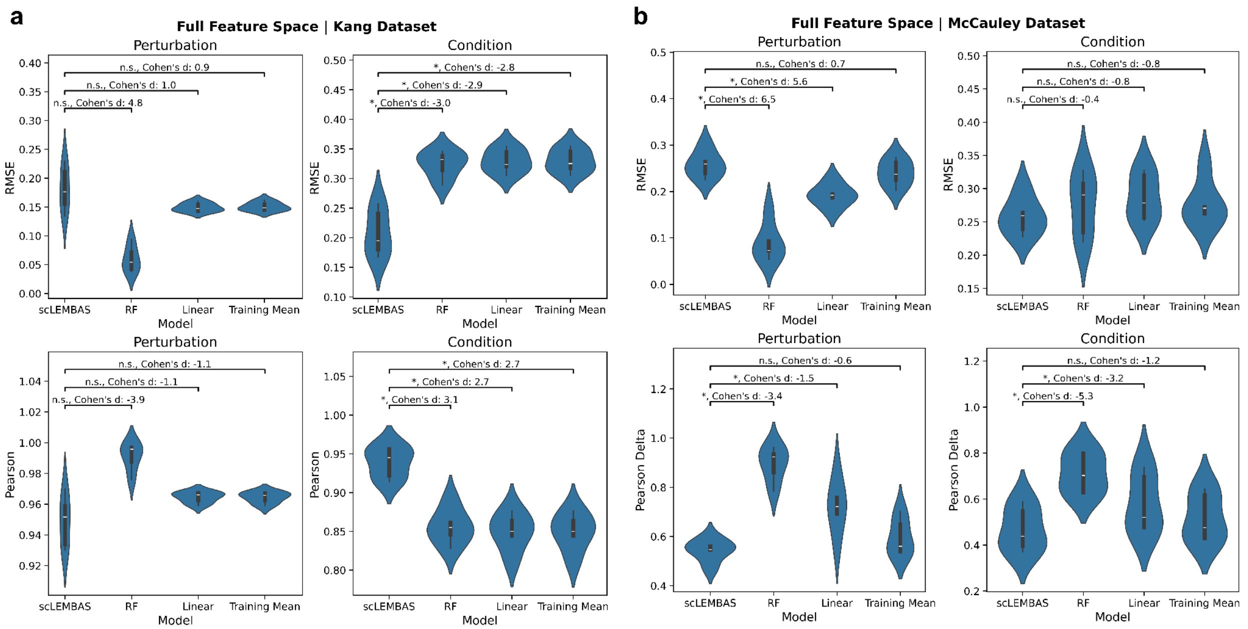


**Figure S9: Pseudobulk baseline quantitative comparisons in full feature space.** For **(a)** the Kang dataset and **(b)** the McCauley dataset: Violin plots show the distribution of test–prediction performance across 5-fold cross-validation, measured by RMSE (top panels) and Pearson delta (bottom panels). Performance is reported for scLEMBAS and baseline models (x-axis), evaluated after pseudo-bulking either by perturbation (left panels) or by cell type–perturbation combinations (right panels). Comparisons are performed between scLEMBAS and each baseline model. Statistical significance is indicated using two-sided Wilcoxon signed-rank tests with Benjamini–Hochberg FDR correction (*, q≤1e-1; **, q≤1e-2).

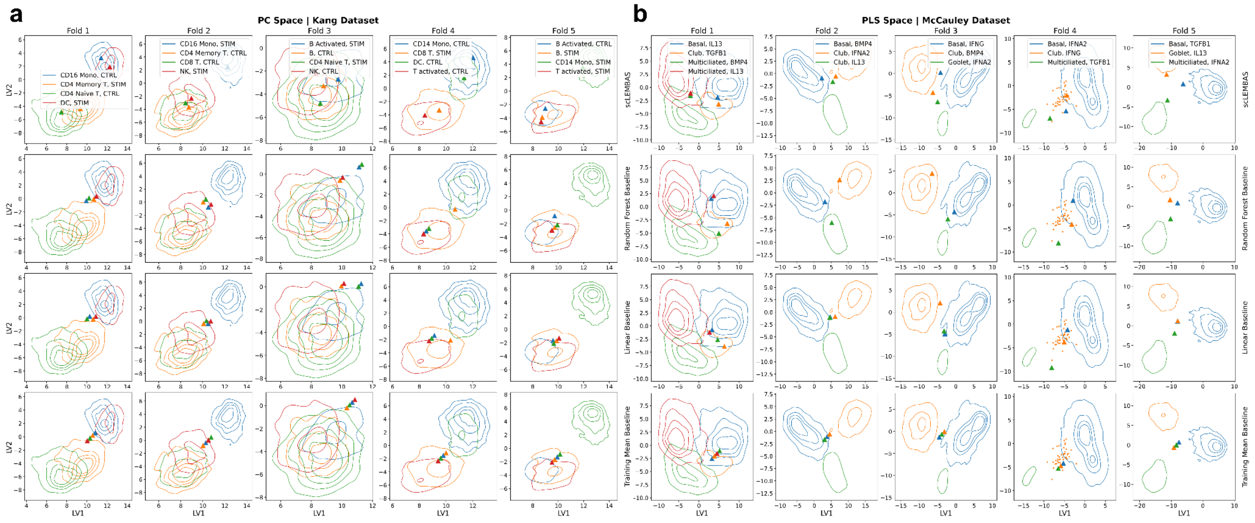


**Figure S10: Pseudobulk baseline visualizations in latent space.** For **(a)** the Kang dataset and **(b)** the McCauley dataset: For each fold, the first two latent variables from models fit on the actual data are shown. For the McCauley dataset, the reduced latent space is defined using PLS with the test conditions as the response variable. For the Kang dataset, the latent space is defined using PCA fit on the entire dataset. Test data are visualized as two-dimensional kernel density estimate (KDE) plots when more than 50 cells are available, and as scatter points otherwise. Predicted condition-level pseudo-bulked outputs are shown as scatter triangles. When model predictions are identical across conditions (e.g., repeated perturbation-level predictions for the RF and linear baselines or across all conditions for the training-mean baseline), small jitter is added for visual clarity.

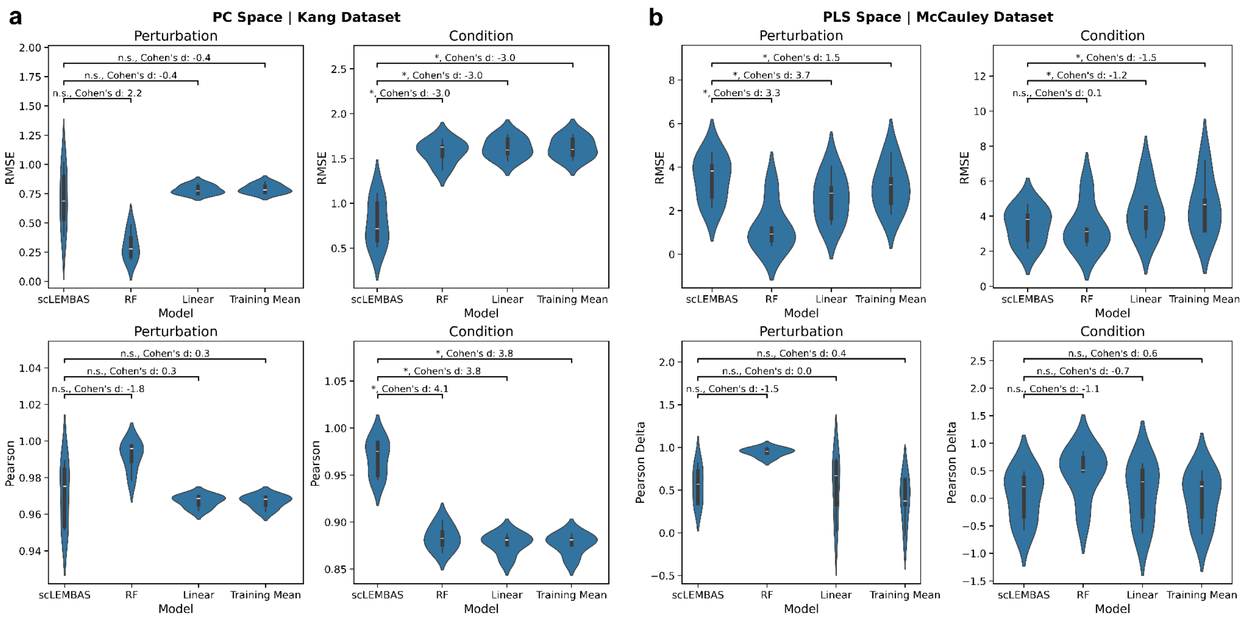


**Figure S11: Pseudobulk baseline quantitative comparisons in reduced feature space.** For **(a)** the Kang dataset and **(b)** the McCauley dataset: Same as Fig. A_E, but evaluated in the respective reduced latent space for each dataset, rather than the full feature space. For the McCauley dataset, the reduced latent space is defined using PLS with the test conditions as the response variable. For the Kang dataset, the latent space is defined using PCA fit on the entire dataset.

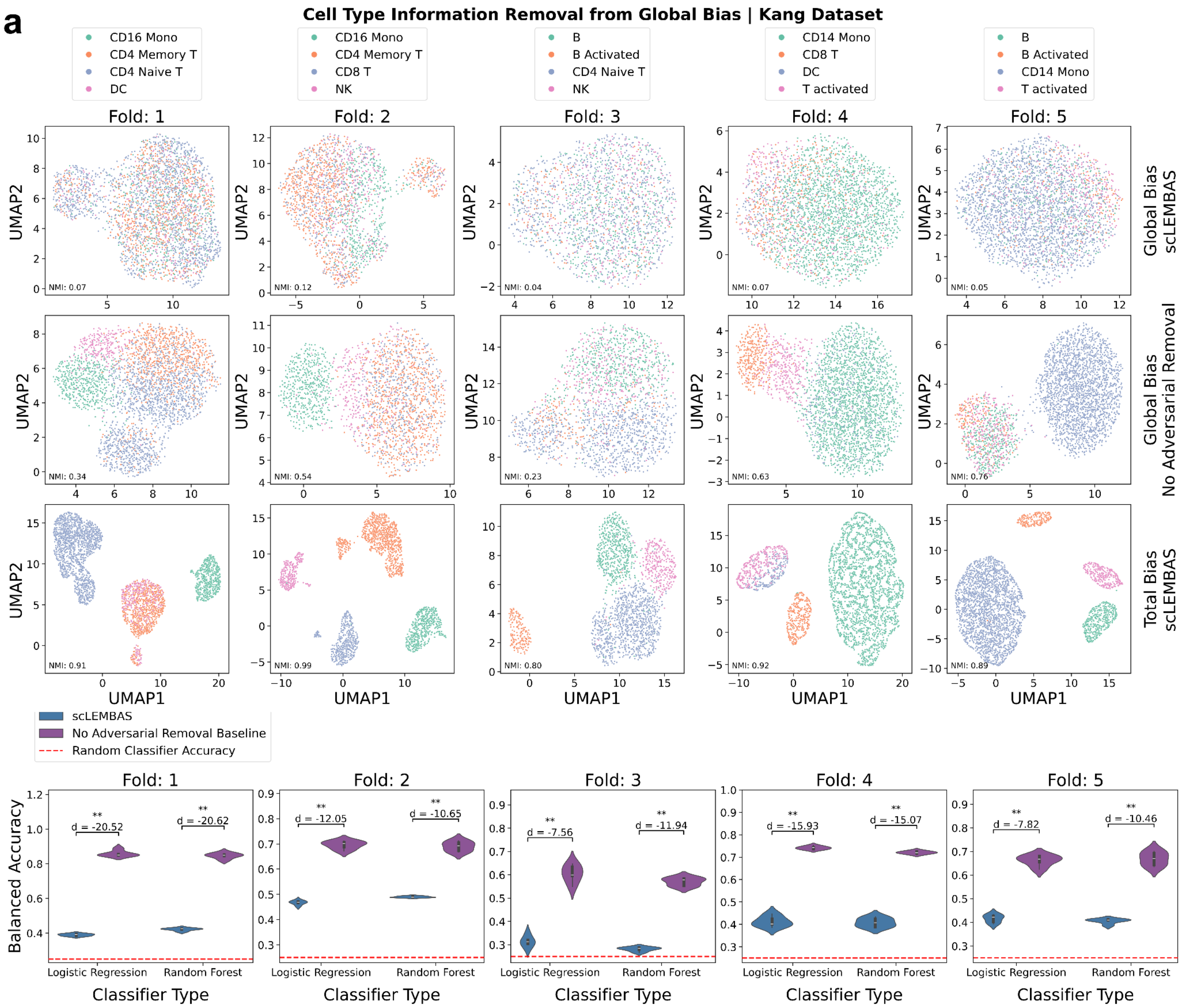
a
Cell Type Information Removal from Global Bias | Kang Dataset
Fold: 1
Fold: 2
Fold: 3
Fold: 4
Fold: 5
Global Bias scLEMBAS
Global Bias No Adversarial Removal
Total Bias scLEMBAS
UMAP1
UMAP2
scLEMBAS
No Adversarial Removal Baseline
Random Classifier Accuracy
Balanced Accuracy
Classifier Type
Logistic Regression
Random Forest

**b**

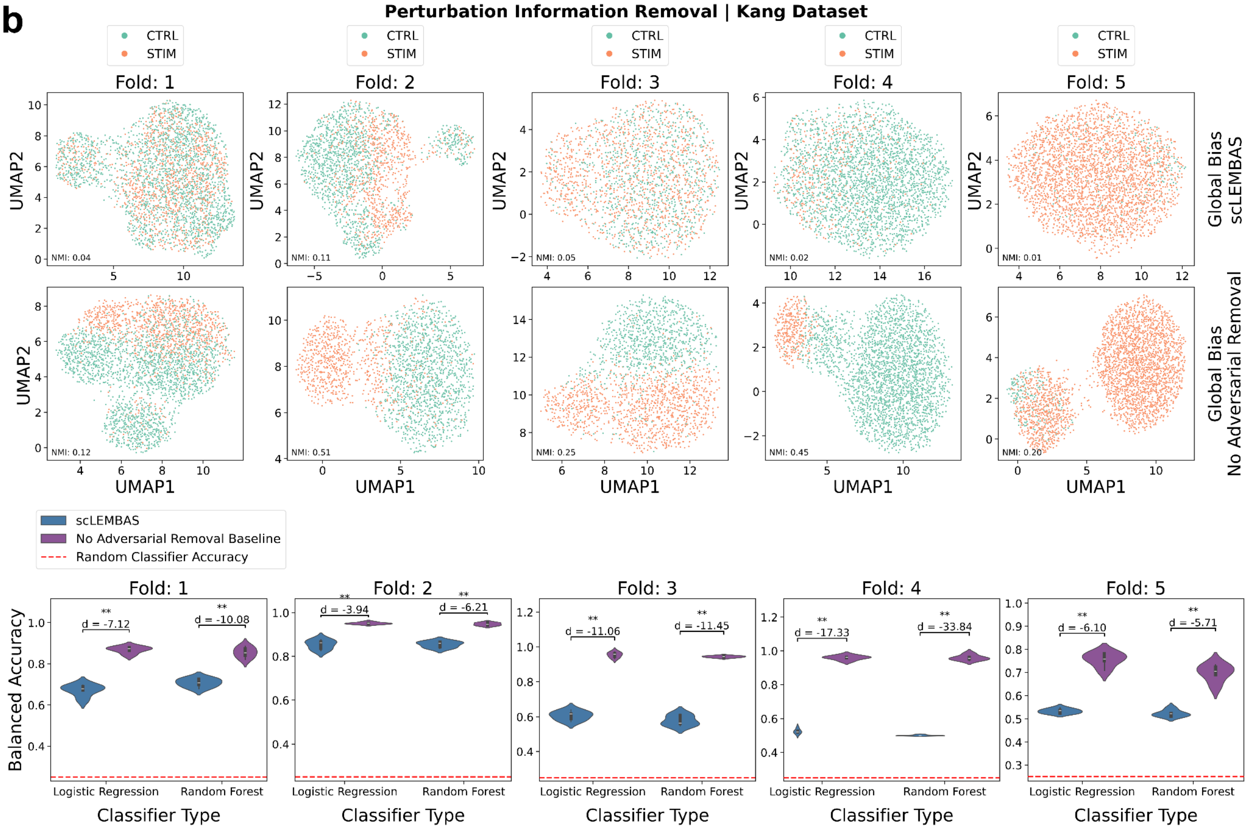
Perturbation Information Removal | Kang Dataset
CTRL
STIM
Fold: 1
Fold: 2
Fold: 3
Fold: 4
Fold: 5
UMAP1
UMAP2
NMI: 0.04
NMI: 0.11
NMI: 0.05
NMI: 0.02
NMI: 0.01
NMI: 0.12
NMI: 0.51
NMI: 0.25
NMI: 0.45
NMI: 0.20
Global Bias scLEMBAS
Global Bias No Adversarial Removal
scLEMBAS
No Adversarial Removal Baseline
Random Classifier Accuracy
Balanced Accuracy
Classifier Type
Logistic Regression
Random Forest
d = -7.12
d = -10.08
d = -3.94
d = -6.21
d = -11.06
d = -11.45
d = -17.33
d = -33.84
d = -6.10
d = -5.71

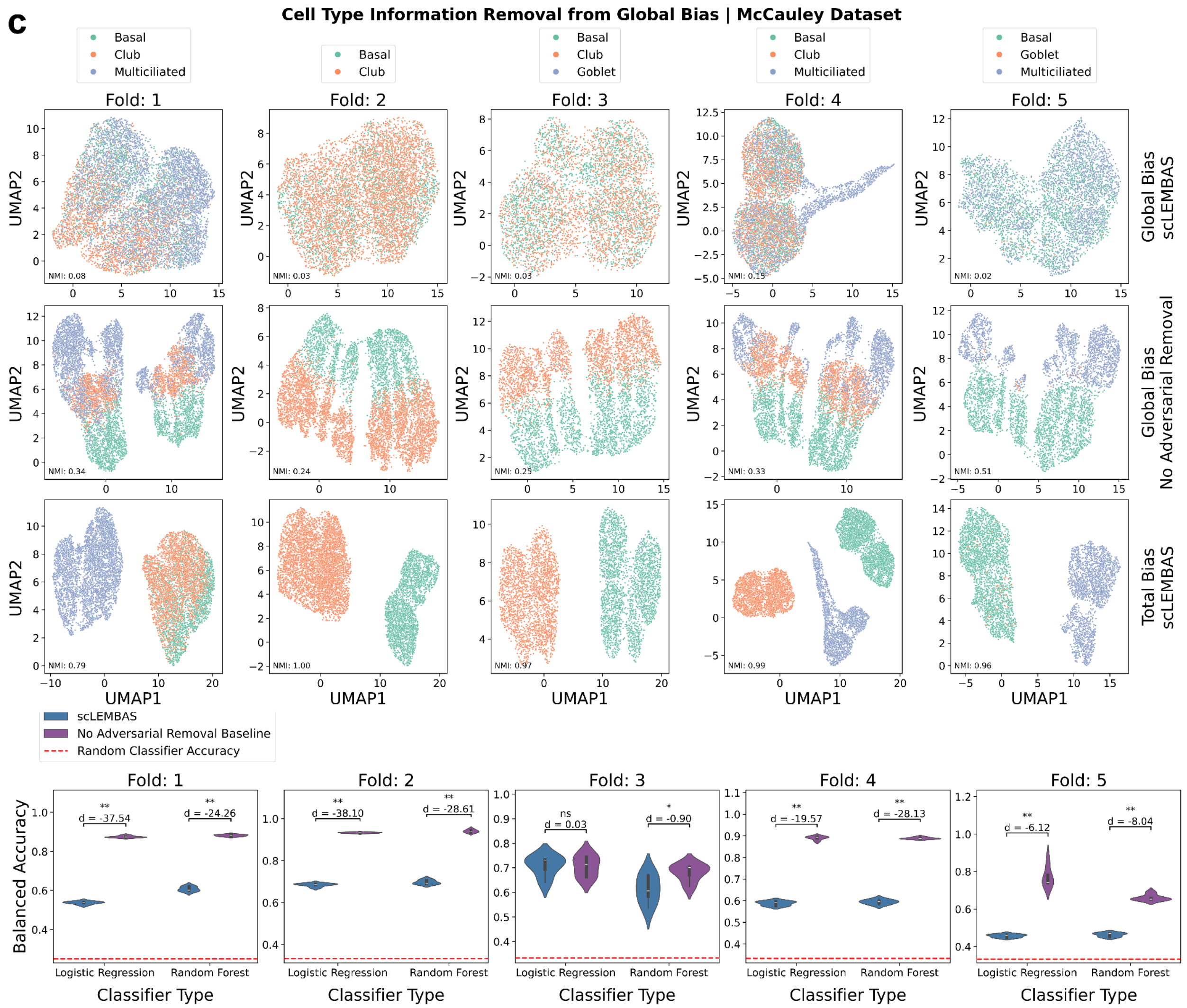


**Figure S12: Covariate information removal from global bias**. Here, we show results for **(a)** cell type information removal from the McCauley dataset, **(b)** cell type removal information removal from the Kang dataset, and **(c)** perturbation information removal from the Kang dataset. The first three rows show UMAPs of test conditions colored by perturbation or cell type. The top row shows global bias generated from the standard scLEMBAS model in each OOD test condition. The second row shows the global bias from the baseline model without adversarial penalization. The third row shows the total bias (global bias + categorical bias) from the standard scLEMBAS model. Panels are annotated with the normalized mutual information between the label of interest and the Leiden cluster. The fourth row shows violin plots for a given test split comparing probe classifier 5-fold CV balanced accuracy between the global bias output by scLEMBAS and the baseline model without adversarial removal. Statistical significance is indicated by the Benjamini–Hochberg FDR-corrected one-sided Mann–Whitney U test (* $q \leq 0.05$, ** $q \leq 0.01$). Probe classifiers were trained to predict the target label using features from each bias's principal component space. The red dashed line indicates the expected balanced accuracy of a random classifier. The McCauley dataset does not show perturbation information removal because all inputs in this standard 5-fold CV were from the same perturbation condition (control, unperturbed cells). The perturbation information removal from the Kang dataset does not show the total bias since this is just addition of cell type information, which is not relevant for this case.

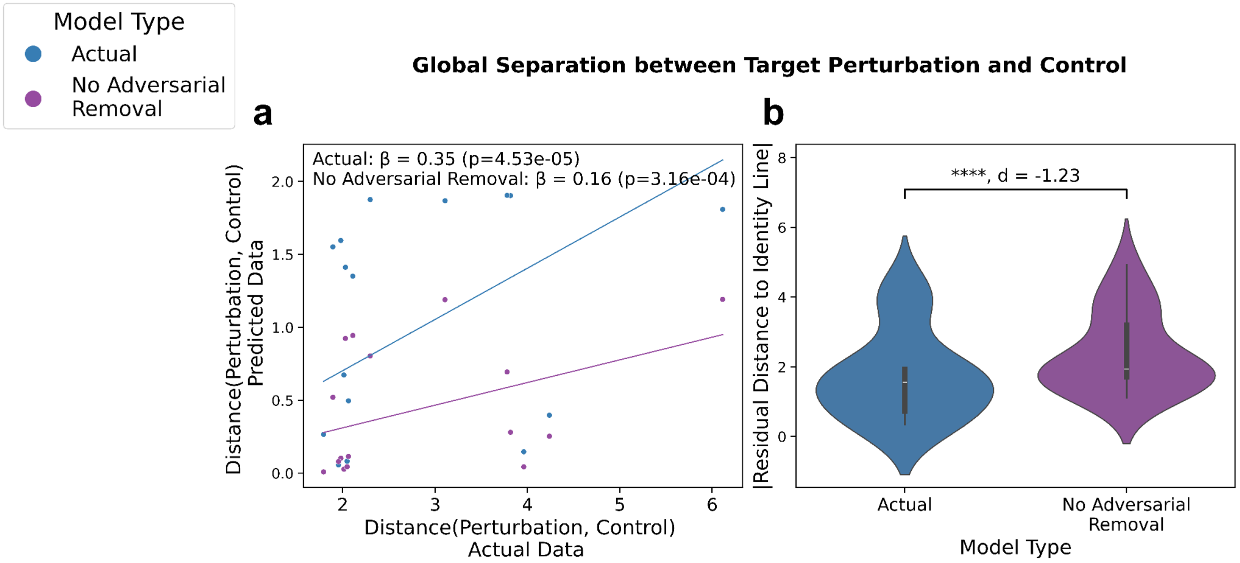


**Figure S13: Global quantification of prediction separation between counterfactual and control. (a)** Scatter plots and OLS regression lines compare mahalanobis distances derived from predicted and actual data. Each point corresponds to a single test condition aggregated across the five cross-validation folds. Mahalanobis distances were computed in PLS space a perturbation and its respective (same cell type) control for each test condition (with the response as perturbation), quantifying counterfactual separation. The predicted distance was between the test split perturbation predicted by the standard scLEMBAS forward pass and the corresponding train split control (same cell type, no perturbation) predicted by the scLEMBAS forward pass without a counterfactual. The actual distance was simply these same two conditions in the actual data. OLS coefficients are annotated alongside their Wald test p-values. Coefficients were estimated using models without an intercept to enable direct comparison to the identity line (gray dashed line), with coefficients closer to 1 indicating more accurate preservation of perturbation separation. **(b)** Violin plots summarize the absolute distance of each scatter point to the identity line, representing deviation from perfect prediction. Comparisons between scLEMBAS and the random baseline model are annotated with a two-sided paired Wilcoxon signed-rank test (* $p \leq 0.05$, ** $p \leq 0.01$, *** $p \leq$ 1e-3, ****, $p \leq$ 1e-4) and a paired Cohen's d effect size.

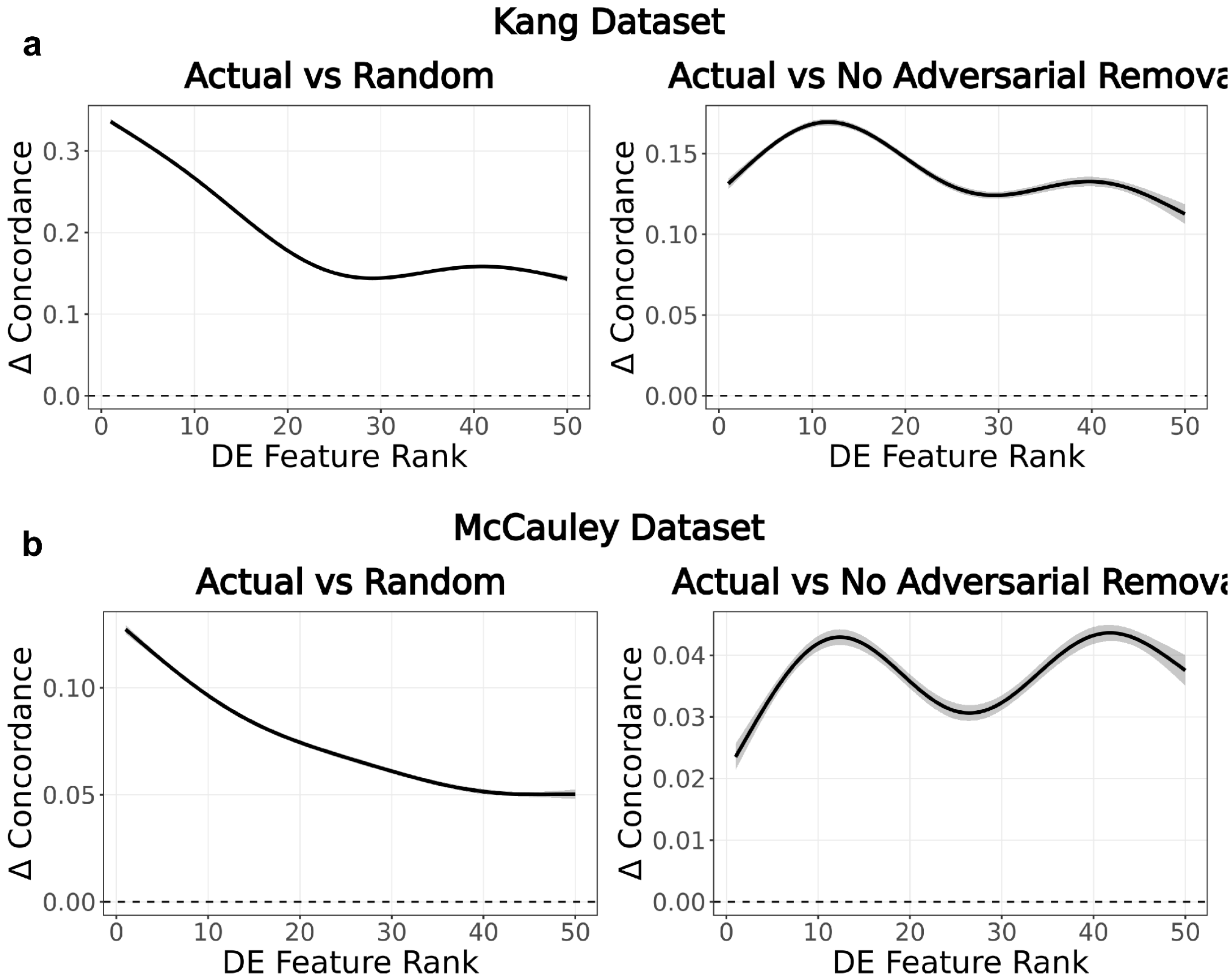


**Figure S14: scLEMBAS' concordance improvement over baselines.** For **(a)** the Kang dataset and **(b)** the McCauley dataset: Difference in GAMM-fitted differential expression concordance between scLEMBAS and baseline models at each DE rank. Positive differences indicate that the actual model outperforms the baselines.

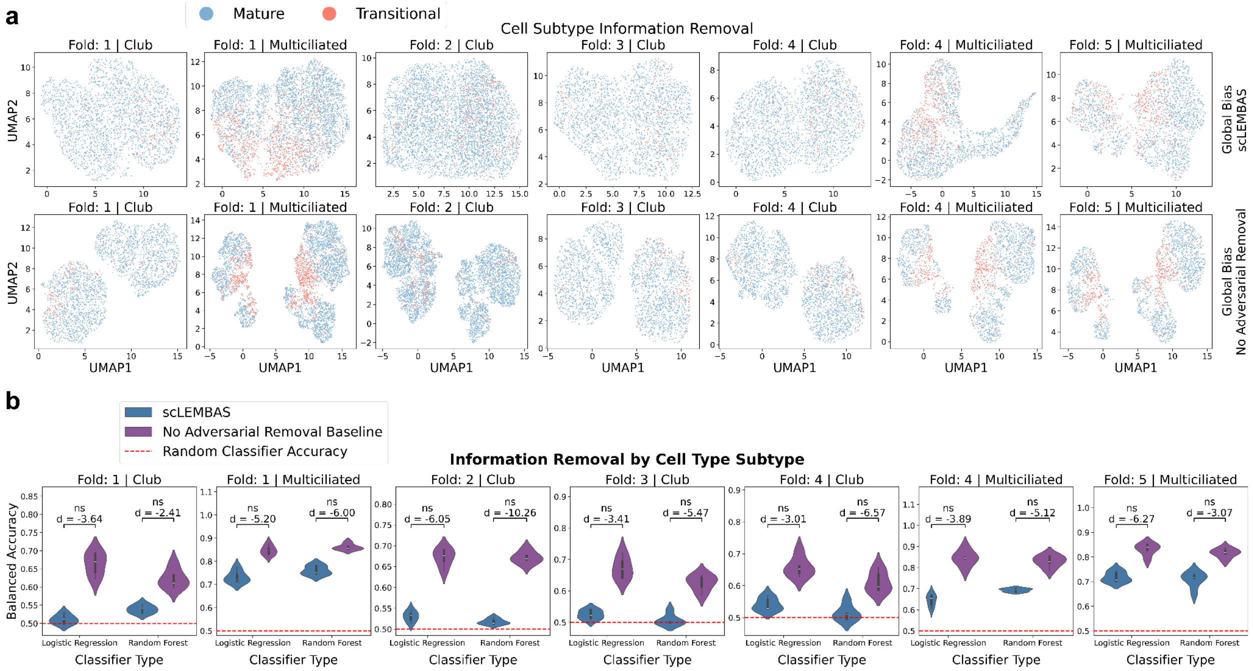


**Figure S15: Cell subtype information in global bias is retained despite adversarial training. (a)** UMAPs using the standard PCA and UMAP embedding pipeline on generated bias terms, stratified by cell type, are colored by cell subtype. UMAPS are displayed for the global bias generated by scLEMBAS (top row) and the global bias generated by the baseline model without adversarial penalization (bottom row). **(b)** Violin plots comparing probe classifier 5-fold cross-validated balanced accuracy between the global bias output on OOD test conditions by scLEMBAS and the baseline model without adversarial removal. Statistical significance is indicated by the Benjamini–Hochberg FDR-corrected one-sided Mann–Whitney U test (* q ≤ 0.05, ** q ≤ 0.01). Probe classifiers were trained to predict the target cell subtype label using features from each model's principal component space. The red dashed line indicates the expected balanced accuracy of a random classifier. In contrast to the analysis on cell type and perturbation labels, cell subtypes were first stratified by cell type to avoid confounding due to dependencies between the two labels. This is analogous to Fig. S12 for cell types and perturbation labels.

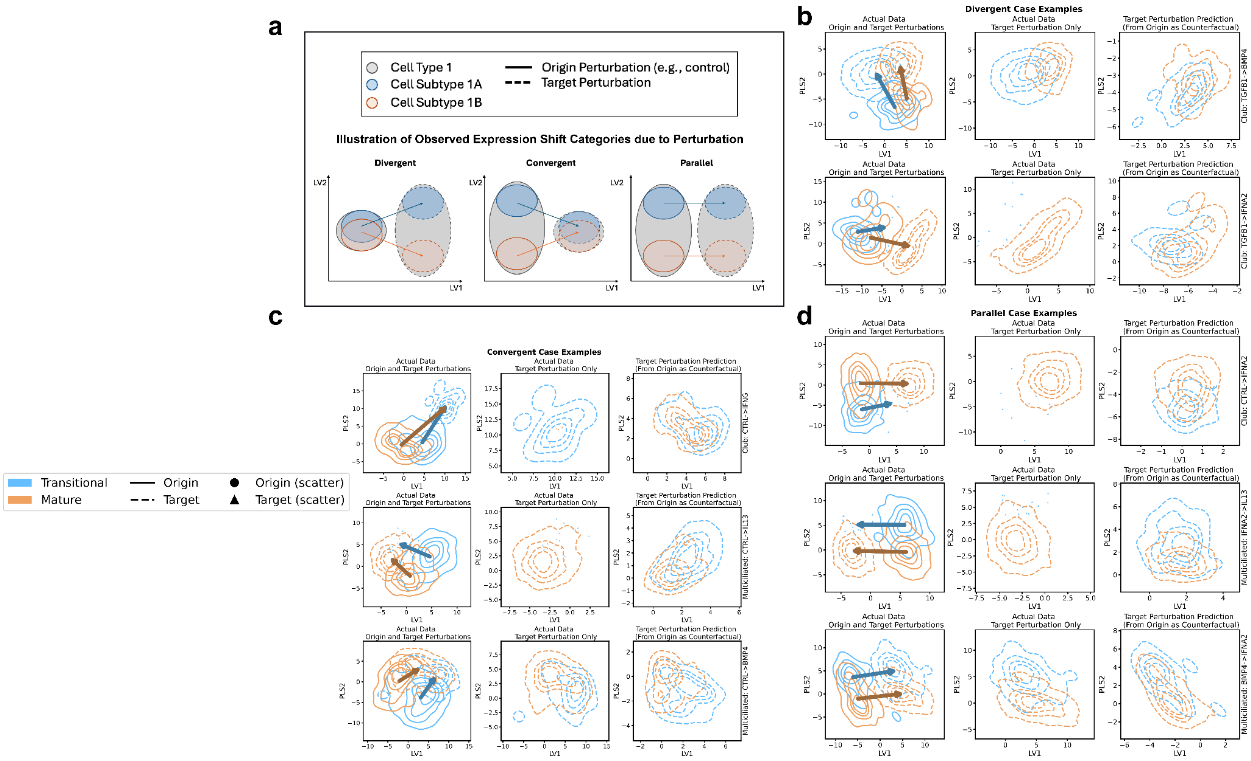


**Figure S16: Three categories of subtype expression shift geometries in response to perturbation.** **(a)** Toy examples demonstrate the three categories of expression shifts induced by transitioning from a starting (i.e., origin) perturbation to a destination (i.e., target) perturbation. Axes represent a theoretical feature space (e.g., latent dimensions). Arrows denote expression shifts, originating at the subtype-specific centroid under the origin perturbation and terminating at the corresponding centroid under the target perturbation. scLEMBAS' counterfactual predictions take as input gene expression from cells under the origin perturbation and predict their TF activity under the target perturbation. Case examples of expression shifts for **(b)** divergent, **(c)** convergent, and **(d)** parallel geometries. For each perturbation and cell subtype pairs, the PLS-reduced space is defined using each of the perturbation and cell subtype labels as a multivariate response, and the first two latent variables are shown. Data are visualized as two-dimensional kernel density estimate (KDE) plots when more than 25 cells are available, and as scatter points otherwise. PLS was fitted using the actual data, after which predictions were projected into the learned latent space. The left panels visualized the actual data for both predicted and target perturbations. Arrows denote expression shifts, originating at the subtype-specific centroid under the origin perturbation and terminating at the corresponding centroid under the target perturbation. The middle panels show only the target perturbation in the actual data, enabling direct visual comparison to the relative geometry of predictions in the right panel. The right hand side is labeled with the cell type, the origin perturbation, and the target perturbation.

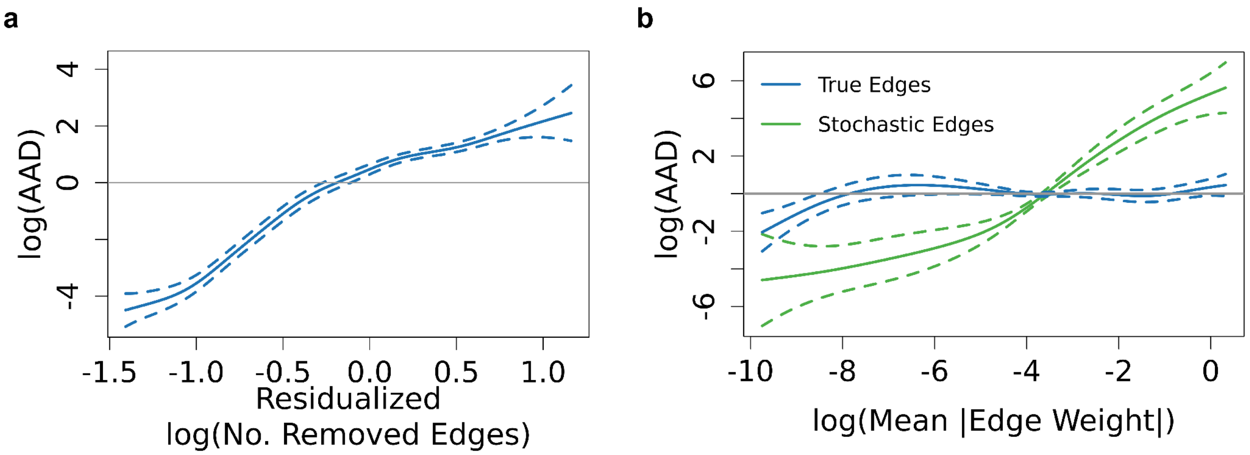


**Fig S17**: **Additional partial effects of scLEMBAS edge removal on AAD**. **(a)** Fitted partial effects from the GAMM showing the association of AAD with the residualized number of edges removed. **(b)** Fitted partial effects from the GAMM that includes the interaction term between edge weight and edge type showing the association of AAD with edge weight stratified by edge type.

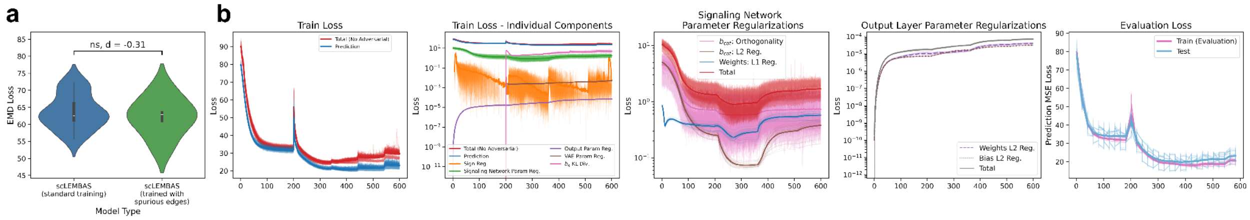


**Fig. S18: Training results and dynamics for ensemble models with spurious interactions added.** Models with L1 regularization are trained on the McCauley dataset using the same 5-fold CV as the standard model split. **(a)** Violin plots comparing EMD test loss for the ensemble models to that of the 5-fold CV in the standard training set up. Ensemble models are trained with an L1 regularization coefficient of 1e-3, whereas the standard models are trained with an L2 regularization coefficient of 1e-7. The panel is annotated with the Cohen's *d* effect size and the two-sided Mann-Whitney U p-value (n.s., $p > 0.05$). **(b)** Training loss and regularization values across epochs for ensemble models. The bold line displays the average across models, whereas the lighter lines display results for each individual model. These panels are analogous to those shown for the standard models using L2 regularization in Fig. S7b.

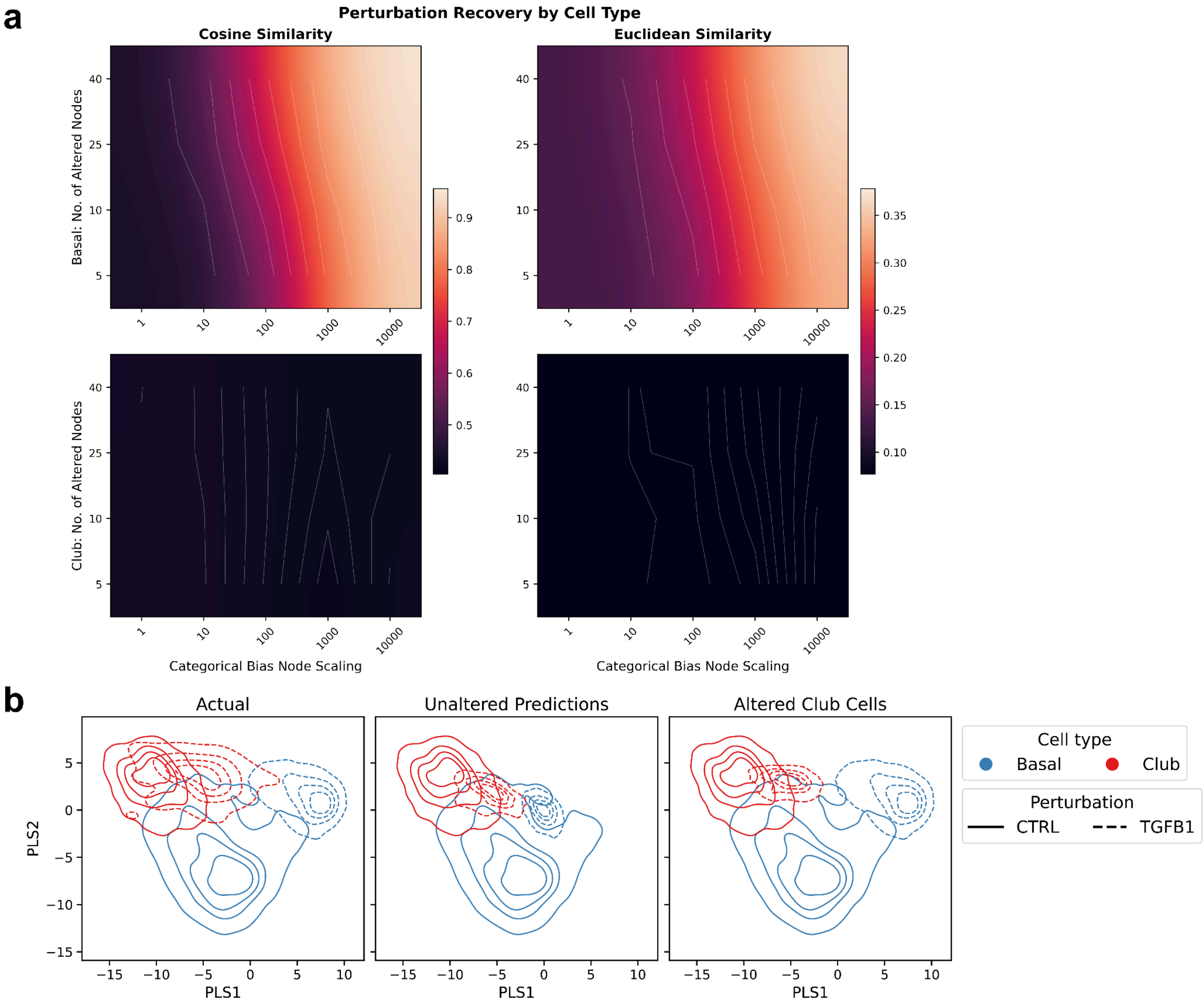


**Fig. S19: (a)** Heatmap of cell-state recovery (left panels - cosine similarity, right panels - Euclidean distance; see Methods for details). Top panels show cell state recovery between predicted basal cells under TGFB1 with altered categorical bias node values and actual TGFB1-perturbed club cells; bottom panels show predicted club cells under TGFB1 with altered categorical bias node values and actual TGFB1-perturbed basal cells. Alterations were conducted across a grid of perturbation parameters: the number of altered nodes (y-axis) and the node scaling factor (x-axis). Results are smoothed via bicubic interpolation and Gaussian filtering. **(b)** Two-dimensional kernel density estimate (KDE) plots of club and basal cells in the first two latent variables of the PLS-reduced space. Solid lines represent unperturbed cell types, and dashed lines represent TGFB1-perturbed cell types. Unperturbed cell types always correspond to the actual data. PLS was fitted using the actual data, after which predictions were projected into the learned latent space. Plots visualize the actual data (left panel), the model predicted TGFB1 perturbed cell states (middle panel), and the model predictions upon categorical bias alteration to club cells, with TGFB1-perturbed basal cells displayed for the actual data (right panel). For categorical bias alterations, select parameters from the parameter sweep were chosen for representative visualization: the scaling factor was set to 1e4, using the top 40 candidate proteins. This is analogous to Fig. 4d, but the right panel shows the alterations to club cells rather than basal cells.

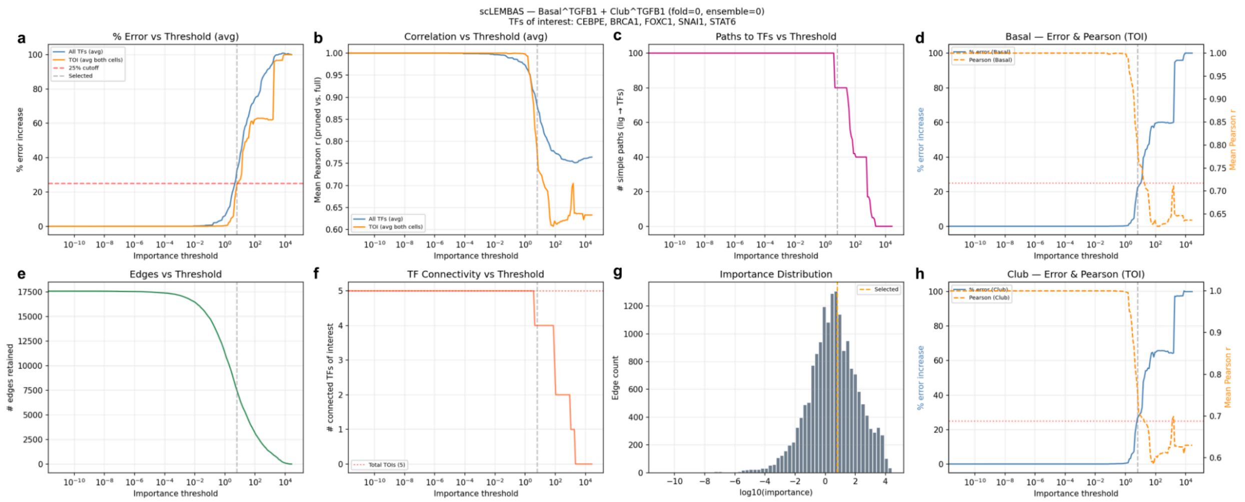


**Figure S20: Per-model diagnostics for importance-threshold selection during TGFB1 subnetwork extraction.** The example shown corresponds to ensemble model 0 from cross-validation fold 0, with CEBPE, BRCA1, FOXC1, SNAI1, and STAT6 specified as transcription factors (TFs) of interest (TOIs). For each candidate importance threshold, signaling edges below the threshold were removed and the resulting unit-TGFB1 counterfactual predictions were compared with those of the complete model. Graph-based connectivity and path metrics were used as diagnostics and did not constrain threshold selection. These metrics were calculated before receptor selection and the final restriction of the subnetwork to directed TGFB1-to-TOI paths. (**a**) Normalized mean absolute error (MAE) across all modeled TF outputs (blue) or the five TOIs (orange). Error was normalized to the corresponding MAE obtained when all signaling network weights were set to zero and is presented as a percentage. TOI error was calculated separately for basal and club cells and then averaged equally across the two cell types. The selected threshold (vertical gray dashed line) was the largest threshold for which the average TOI error did not exceed 25% (horizontal red dashed line). (**b**) Mean TF-wise Pearson correlation between the predictions of the thresholded and complete models, calculated across cells and averaged across basal and club cells for either all modeled TFs (blue) or the TOIs (orange). TFs with invariant predictions in either model were excluded from the correlation calculation. (**c**) Number of simple directed paths connecting TGFB1 to the TOIs represented in the signaling graph. Paths were restricted to a maximum of five edges and capped at 20 paths per TOI. (**d**) Basal cell fidelity across thresholds, quantified using normalized TOI error (blue; left axis) and mean TF-wise Pearson correlation (orange dashed; right axis). (**e**) Number of signaling edges retained at each importance threshold. (**f**) Number of TOIs reachable from TGFB1 in the thresholded graph; the horizontal red dotted line indicates the total number of TOIs represented as signaling nodes. (**g**) Distribution of positive edge-importance scores on a $log_{10}$ scale. The orange dashed line indicates the selected threshold. (**h**) Club cell normalized TOI error and mean TF-wise Pearson correlation, defined as in panel D. The 25% lines in panels D and H are provided as cell-type-specific references; threshold selection was based on the TOI error averaged across both cell types in panel A.
